# Contribution of circulating host and microbial tryptophan metabolites towards Ah receptor activation

**DOI:** 10.1101/2023.01.26.525691

**Authors:** Ethan W. Morgan, Fangcong Dong, Andrew Annalora, Iain A. Murray, Trenton Wolfe, Reece Erickson, Krishne Gowda, Shantu G. Amin, Kristina S. Petersen, Penny M. Kris-Etherton, Craig Marcus, Seth T. Walk, Andrew D. Patterson, Gary H. Perdew

## Abstract

The aryl hydrocarbon receptor (AHR) is a ligand activated transcription factor that plays an integral role in homeostatic maintenance by regulating cellular functions such as cellular differentiation, metabolism, barrier function, and immune response. An important but poorly understood class of AHR activators are compounds derived from host and bacterial metabolism of tryptophan. The commensal bacteria of the gut microbiome are major producers of tryptophan metabolites known to activate the AHR, while the host also produces AHR activators through tryptophan metabolism. We used targeted mass spectrometry-based metabolite profiling to determine the presence and metabolic source of these metabolites in the sera of conventional mice, germ-free mice, and humans. Surprisingly, sera concentrations of many tryptophan metabolites are comparable between germ-free and conventional mice. Therefore, many major AHR-activating tryptophan metabolites in mouse sera are produced by the host, despite their presence in feces and mouse cecal contents. AHR activation is rarely studied in the context of a mixture at relevant concentrations, as we present here. The AHR activation potentials of individual and pooled metabolites were explored using cell-based assays, while ligand binding competition assays and ligand docking simulations were used to assess the detected metabolites as AHR agonists. The physiological and biomedical relevance of the identified metabolites was investigated in the context of cell-based models for cancer and rheumatoid arthritis. We present data here that reframe AHR biology to include the presence of ubiquitous tryptophan metabolites, improving our understanding of homeostatic AHR activity and models of AHR-linked diseases.

## Introduction

Maintaining homeostasis is fundamental to life. Homeostasis is a broad term that includes the many cellular activities that facilitate normal bodily functioning. Evolution has contributed exceptionally complex biochemical and molecular schemes to tightly regulate cellular activity through cell-cell signaling, environmental sensors, transcription factors, post-translational modifications and more. Increased complexity enables greater regulatory control and management of large systems, as seen in mammalian organisms with highly specialized cell types. An important factor in maintaining the specificity of mammalian homeostasis is the ligand activated transcription factor, aryl hydrocarbon receptor (AHR). The AHR, a member of the basic helix-loop-helix/Per-Arnt-Sim family, possesses a promiscuous ligand binding pocket, which allows the recognition of diverse ligands, thus providing flexibility in its function. Upon ligand-mediated activation, the AHR translocates from the cytoplasm to the nucleus where it heterodimerizes with AHR nuclear translocator. This heterodimer binds dioxin response elements in the promoter region of AHR-responsive genes, thereby influencing gene expression.^1^ The genes under the AHR’s influence span a variety of cellular functions, notably these involved in cellular differentiation, xenobiotic metabolism, and immune function.

The AHR is often studied in terms of facilitating the metabolism of toxic environmental pollutants including polycyclic aromatic hydrocarbons like 2,3,7,8-tetrachlorodibenzo-*p*-dioxin (TCDD) or benzo[*α*]pyrene.^1^ It is also known to play an important role mediating immune responses by regulating the production of anti- or pro-inflammatory cytokines.^2,3,4^ Additionally, the AHR is involved in cell differentiation of multiple cell types including T cells, B cells, and goblet cells.^3,5,6,7^ Furthermore, the AHR is widely expressed throughout multiple organ systems and is highly expressed in barrier sites where it plays a role in maintaining epithelial barrier integrity.^2,8^ These expression patterns allow the body to recognize ligands and perform cell-specific roles that contribute to the regulation of homeostasis.

Although it is most studied for its recognition of industrial pollutants, the AHR ligand binding pocket can bind many compounds produced by host, bacterial, and plant metabolism of tryptophan (Trp). An estimated 90% of dietary Trp enters the kynurenine pathway, playing an important role in regulating homeostatic Trp concentrations, while also producing compounds capable of activating the AHR, including kynurenine, kynurenic acid, and xanthurenic acid.^9,10,11^ A much smaller proportion of dietary Trp (~2%) enters the serotonin pathway, although these compounds are not known to directly interact with AHR and do not represent a current significant area of research.^12^ Bacteria in the gut microbiome use an estimated 5% of Trp to produce indole-based derivatives. The trillions of cells comprising the gut microbiome produce numerous Trp metabolites for diverse physiological functions, including energy metabolism, cell signaling, and stress response.^13,14,15^ A subset of these compounds are reported to activate the AHR and are major subjects of research, notably, indole, indole-3-propionic acid (IPA), and indole-3-acetic acid (IAA).^10,14,16,17^ Additionally, IAA and IPA are known in the plant sciences as auxins, a class of plant hormones, and thus may be present in our diets.^18,19^ Trp metabolism provides a robust and consistent pool of AHR ligands that could be contributing to the regulation of systemic AHR activity and, accordingly, are contributors to homeostatic maintenance. We refer to this diverse class of metabolites as pseudo-endogenous due to the difficulty in distinguishing the metabolite progenitor from host, microbiome, or diet. Previously, our group has determined the presence of Trp metabolites in feces and cecal contents of mice that at the concentration present can generate AHR activity.^20^ However, the absorption and circulation of these metabolites is still poorly understood in the context of homeostatic AHR activity. Our work provides new information and clarity to this area of research.

In this study, we used liquid chromatography paired with tandem mass spectrometry (LC-MS/MS) to identify and quantify Trp metabolites circulating in human, mouse, and germ-free mouse sera. We identified metabolites frequently attributed to bacterial metabolism that are, in fact, produced independent of bacteria. Further, by profiling circulating metabolites we found that fecal metabolite concentrations do not serve as an accurate proxy measurement for Trp metabolites that are circulating in a host. We demonstrate that circulating Trp metabolites can generate AHR activity, but do not have a synergistic or additive effect and produce varied outcomes in different cell types. Our work, identifying relevant AHR agonists, provides a basis to recontextualize physiologic AHR activation and may help in examining the link between host metabolism and AHR-linked diseases.

## Results

### Tryptophan metabolite profiling of the sera of conventional and germ-free mice

We initially aimed to build upon our previous work analyzing murine fecal and cecal contents for the presence of AHR activating Trp metabolites by analyzing murine sera to investigate into which Trp metabolites in the host circulation are derived from diet, host metabolism and/or the gut microbiome.^20^ Blood was collected from the portal vein of conventional C567BL/6J mice and sera analyzed by targeted LC-MS, with the exception of indole, which was analyzed by GC-MS. Guided by our previous work, a compilation of 16 Trp metabolites known to elicit AHR activity were targeted for quantification (Table 1). Of the 16 metabolites, nine were detected and quantified through interpolation of standard curves and corrected for recovery and matrix effects (Fig. 1, Table S1). Analysis revealed IPA was the most abundant Trp metabolite with a mean concentration of 8.4 ± 5.69 μM and comprising 50.92% of the total composition of examined metabolites by concentration. IPA represents a singularly large proportion of the total examined metabolites with the remaining metabolites composing substantially smaller proportions: 15.68% indole-3-lactic acid (ILA), 11.87% kynurenine (KYN), 7.38% IAA, 5.99% 5-hydroindole-3-acetic acid, 3.12% kynurenic acid (KA), 2.91% indole-3-carboxaldehyde (I3C), 1.10% indole-3-acrylic acid, and 1.02% 2-oxindole. The remaining seven targeted metabolites were undetectable based on their limits of detection (Table 1). These data present a remarkably different metabolic profile compared to the analysis of murine feces or cecal contents.^20^ Additionally, these metabolites, especially the indole-containing compounds, have been linked to the gut microbiome and bacterial metabolism, indicating that the microbiome may indeed serve as an important source of AHR activators in mouse sera^10,20^

**Table 1.**
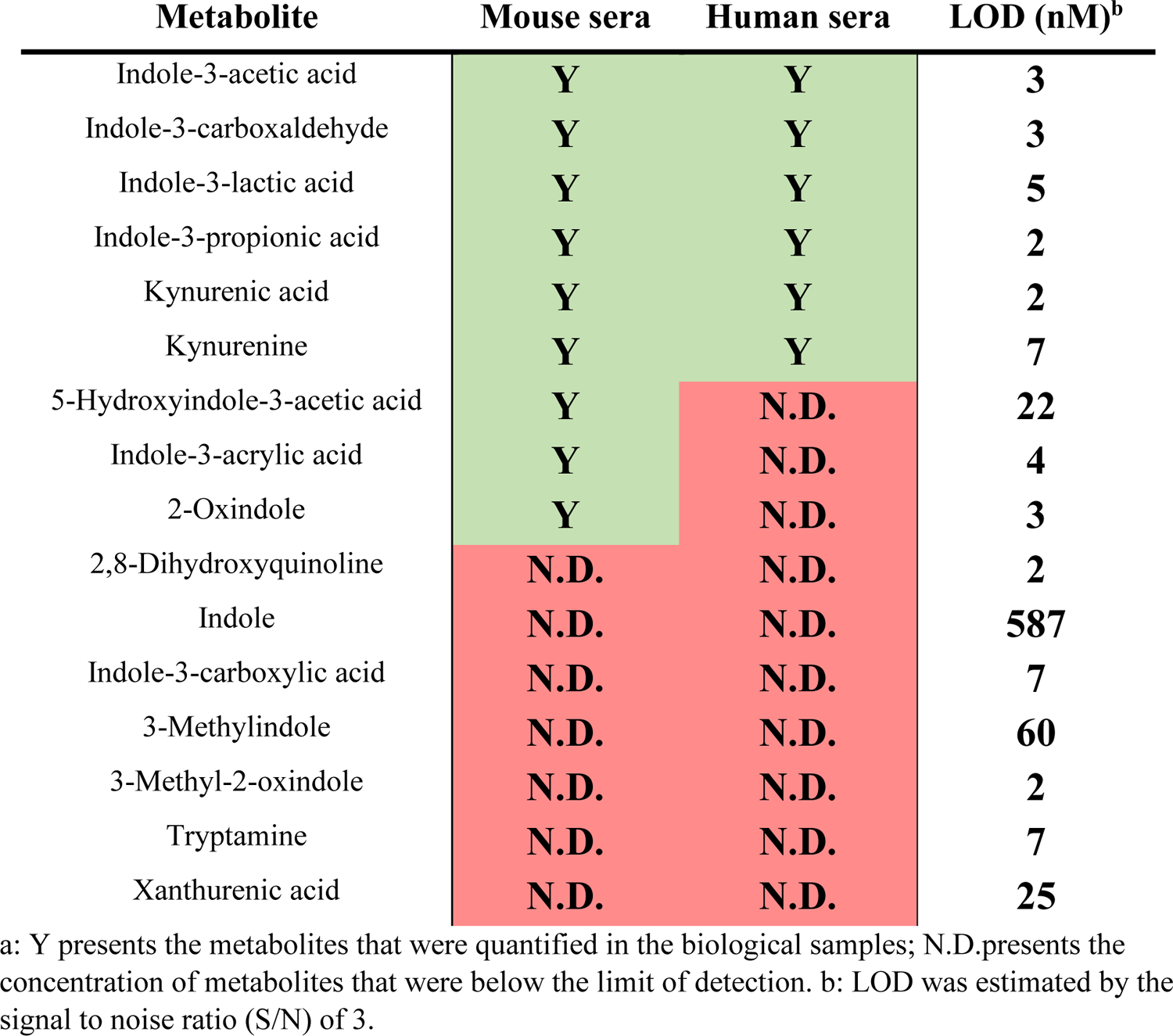
Summary of profiled tryptophan metabolites in mouse serum and human sera^a^.

| **Metabolite** | **Mouse sera** | **Human sera** | **LOD (nM)<sup>b</sup>** |
| --- | --- | --- | --- |
| Indole-3-acetic acid | **Y** | **Y** | **3** |
| Indole-3-carboxaldehyde | **Y** | **Y** | **3** |
| Indole-3-lactic acid | **Y** | **Y** | **5** |
| Indole-3-propionic acid | **Y** | **Y** | **2** |
| Kynurenic acid | **Y** | **Y** | **2** |
| Kynurenine | **Y** | **Y** | **7** |
| 5-Hydroxyindole-3-acetic acid | **Y** | **N.D.** | **22** |
| Indole-3-acrylic acid | **Y** | **N.D.** | **4** |
| 2-Oxindole | **Y** | **N.D.** | **3** |
| 2,8-Dihydroxyquinoline | **N.D.** | **N.D.** | **2** |
| Indole | **N.D.** | **N.D.** | **587** |
| Indole-3-carboxylic acid | **N.D.** | **N.D.** | **7** |
| 3-Methylindole | **N.D.** | **N.D.** | **60** |
| 3-Methyl-2-oxindole | **N.D.** | **N.D.** | **2** |
| Tryptamine | **N.D.** | **N.D.** | **7** |
| Xanthurenic acid | **N.D.** | **N.D.** | **25** |

a: Y presents the metabolites that were quantified in the biological samples; N.D.presents the concentration of metabolites that were below the limit of detection. b: LOD was estimated by the signal to noise ratio (S/N) of 3.

**Figure 1.**
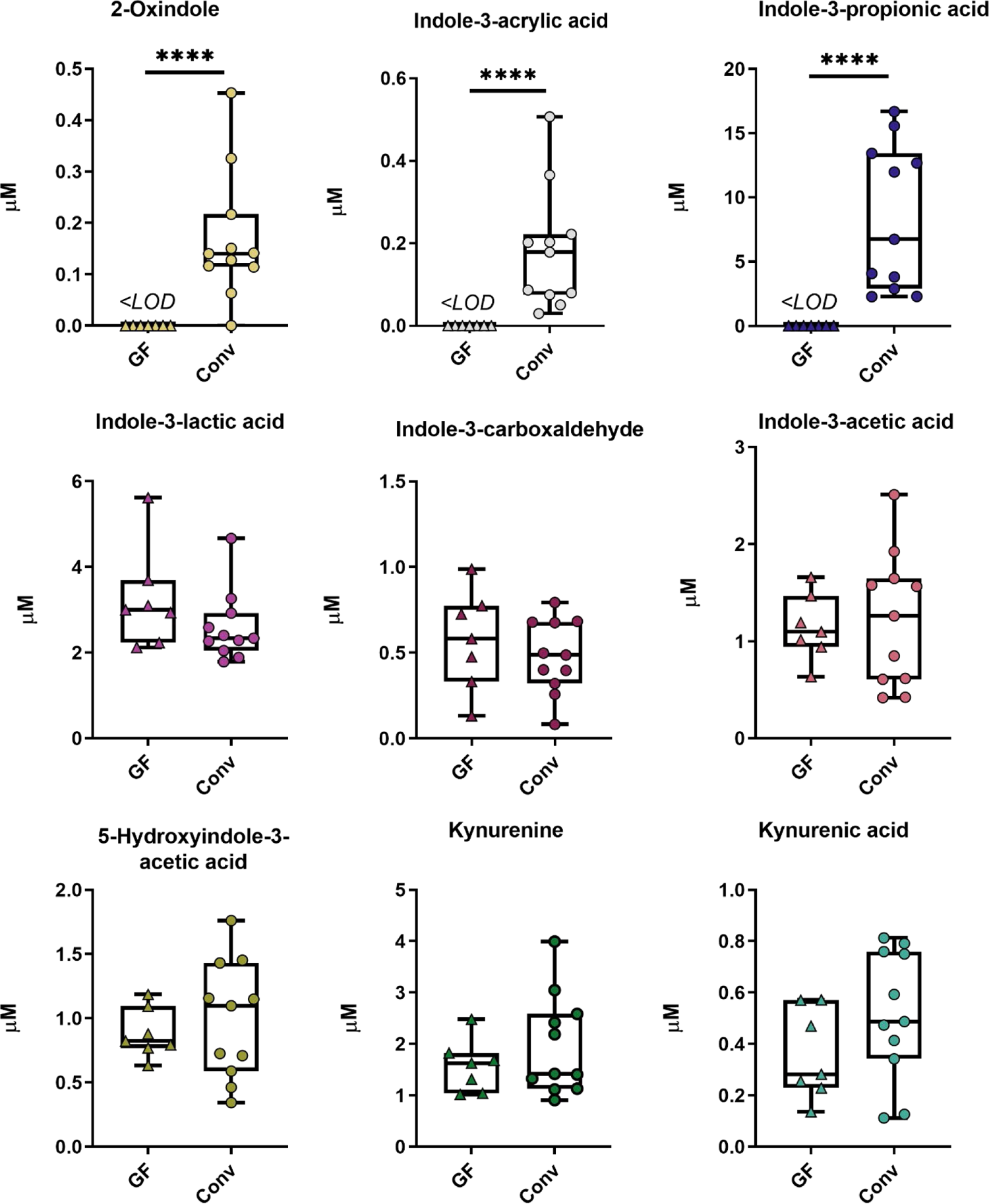
Comparison of tryptophan metabolites in serum of GF mice and conventional mice. LC-MS analysis identified nine Trp metabolites in serum of conventional and GF mice fed a standard chow diet, *ad libitum*. Statistical comparisons were made using Mann-Whitney and unpaired *t-*test. Each box represents the median value with Q1 and Q3 range, and whiskers describe the maximal and minimal values. Conv, conventional mice; GF, germ-free mice; <LOD, below the limit of detection. (***): p-value = 0.0005; (****): p-value < 0.0001.

To test this hypothesis and evaluate the necessity of microorganisms to produce these nine examined metabolites, sera were collected from germ-free (GF) mice and subjected to the same targeted metabolomic analysis. Surprisingly, only IPA, 2-oxindole and indole-3-acrylic acid were undetected in the serum of GF mice. The remaining six metabolites were present and their concentrations were comparable between GF and conventional mice maintained on the same diet (Fig. 1, Table 2). In addition to routine surveillance of GF mice (see Methods), the absence of bacteria e was supported by undetectable serum concentrations of hippuric acid (Fig. S1), a known microbial metabolite.^21^ Taken together, these data indicate that the profile of circulating metabolites contains fewer Trp metabolites than those based on fecal analysis, and indole-containing compounds that are often ascribed to microbial metabolism (e.g. IAA, I3C, ILA) appear to actually originate from host metabolism.

**Table 2.** Comparison of serum Trp metabolite concentrations in conventional mice and GF mice

| **Metabolite** | **Mean abundance ± S.D. (μM)** |  |
| --- | --- | --- |
|  | **Conventional mice** | **GF mice** |
| **5-hydroxyindole-3-acetic acid** | 0.99 ± 0.46 | 0.88 ± 0.19 |
| **Indole-3-acetic acid** | 1.22 ± 0.69 | 1.14 ± 0.34 |
| **Indole-3-acrylic acid** | 0.18 ± 0.15 | <LOD |
| **Indole-3-carboxaldehyde** | 0.48 ± 0.22 | 0.57 ± 0.29 |
| **Indole-3-lactic acid** | 2.59 ± 0.81 | 3.24 ± 1.18 |
| **Indole-3-propionic acid** | 8.4 ± 5.69 | <LOD |
| **Kynurenic acid** | 0.51 ± 0.25 | 0.36 ± 0.18 |
| **Kynurenine** | 1.96 ± 0.97 | 1.57 ± 0.51 |
| **2-Oxindole** | 0.17 ± 0.13 | <LOD |

### Diet alters circulating tryptophan metabolites

Having generated Trp metabolite profiles from sera, we asked if altering the diet would significantly change circulating concentrations. One of two groups of conventional C57BL/6J mice with *ad libitum* access to a standard laboratory chow were switched onto a semi-purified diet (AIN-93G) while the second remained on standard chow. After one week, portal vein blood was collected and analyzed using targeted LC-MS (Fig. 2). The abundance of IPA dropped significantly (39-fold) in mice eating the semi-purified diet with (mean concentration 4.71 ± 1.48 μM and 0.12 ± 0.10 μM for standard and semi-purified diets, respectively). Additionally, the abundance of indole-3-acrylic acid also dropped by 3-fold in the same group of mice (mean concentration 0.31 ± 0.10 μM and 0.10 ± 0.02 μM, respectively). Given that bacteria are necessary for the circulation of IPA and indole-3-acrylic acid (IAcA) in serum (Fig. 1), this result indicates that diet can be used to substantially alter microbial Trp metabolism. Furthermore, except for IAA, the dietary shift had no statistically significant influence Trp metabolite abundances. The difference observed in IAA, while significant, was minor (1.6-fold) compared to that observed in IAcA and IPA. The remaining six targeted metabolites did not significantly differ in abundance. While it is not clear which dietary components influence microbial or host Trp metabolism, the data demonstrate that diet can alter the abundance and distribution of certain circulating Trp metabolites.

**Figure 2.**
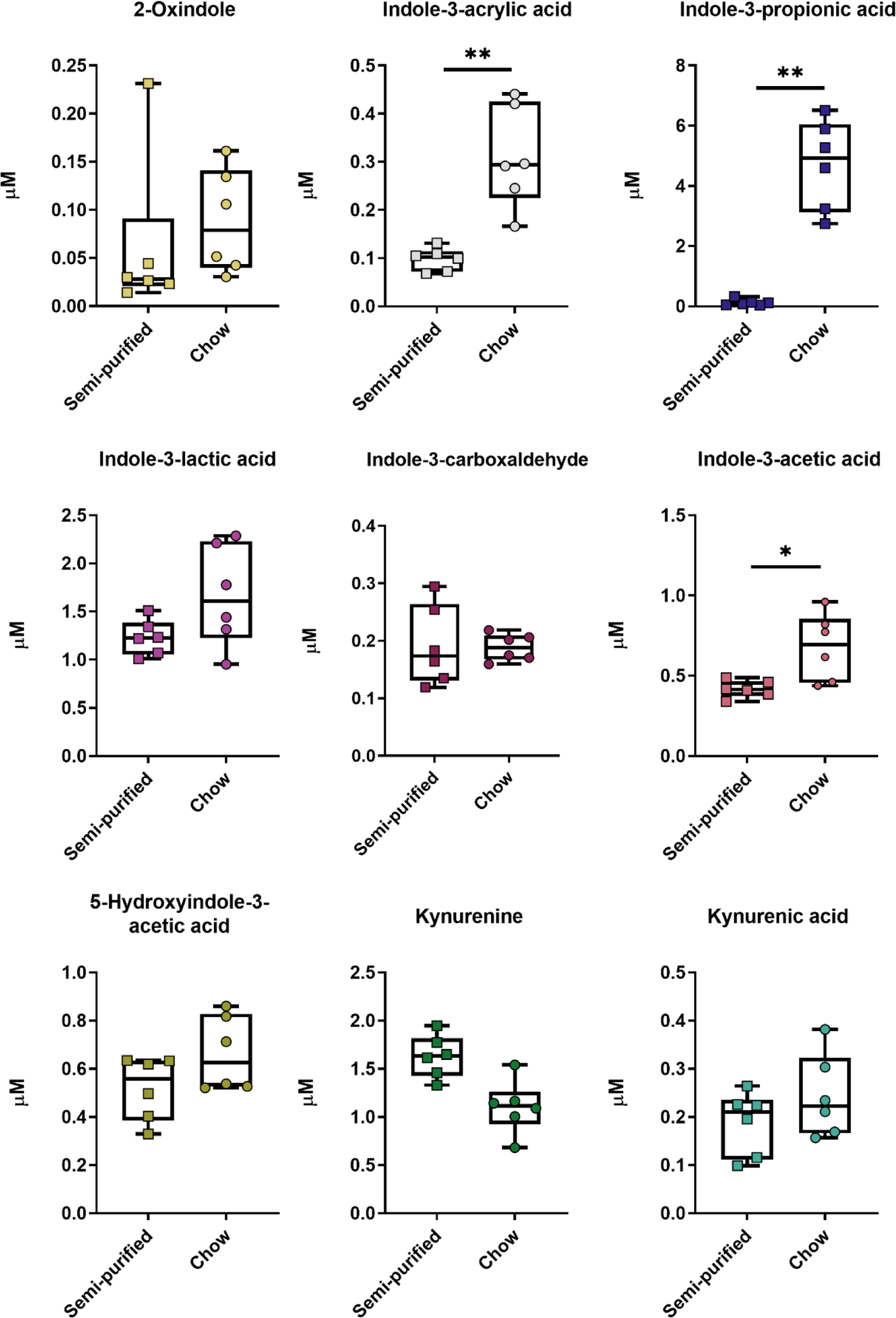
Comparison of tryptophan metabolites in serum of conventional mice fed a standard chow or semi-purified diet. Conventional mice were fed a standard chow diet or semi-purified diet (AIN-93G), *ad libitum*, and the nine Trp metabolites were quantified by LC-MS analysis. Statistical comparisons were made using Mann-Whitney or unpaired *t-*test. Each box represents the median value with Q1 and Q3 range, and whiskers describe the maximal and minimal values. (***): p-value = 0.0005; (****): p-value < 0.0001.

### Activation of murine AHR by circulating tryptophan metabolites

As we and others have observed, the detected metabolites possess the potential to activate the AHR, so we next assessed the ability for these nine metabolites to produce AHR activity at their relevant serum concentrations.^20,22,23,24^ We assessed the potential of each metabolite to induce AHR activity at the mean concentrations measured in mouse sera (Table 3). We treated a murine AHR-dependent reporter assay system (Hepa 1.1) to each of the nine metabolites for 4 h. We also generated a synthetic Trp metabolite pool using the mean concentrations of all nine quantified Trp metabolites to model the potential for the complete Trp metabolite profile to generate AHR activity (Fig. 3). The complete Pool generated a ~7-fold induction (p < 0.0001) relative to the vehicle treatment. KYN was the only individual metabolite to produce an increase in luciferase activity (~6-fold induction, p <0.0001) relative to the vehicle treatment. Despite KYN comprising just 12% of the total concentration of the Pool, it generated 85% the activity produced by the whole Pool.

**Figure 3.**
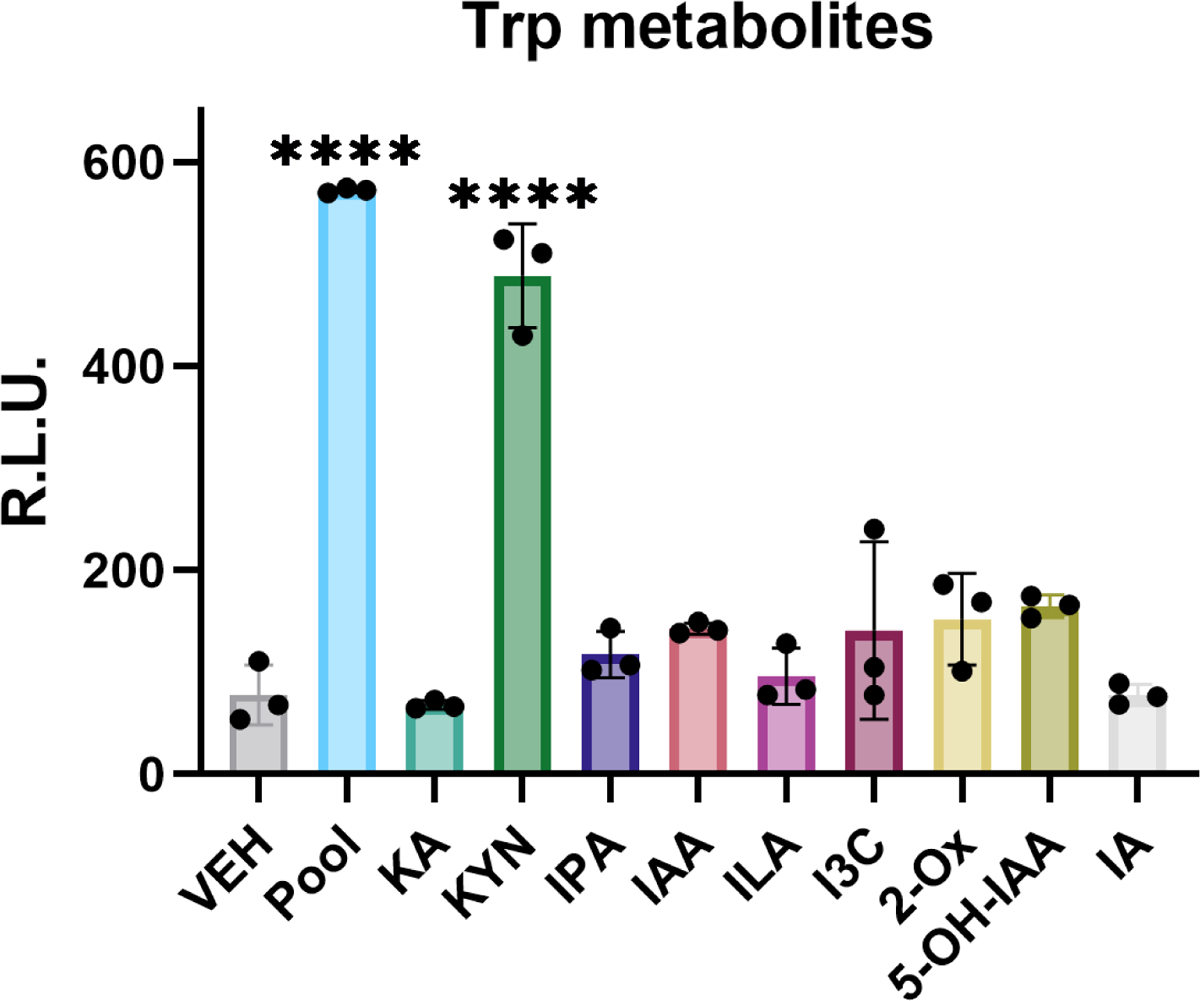
Mouse AHR activity generated from circulating Trp metabolites. Mean concentrations of Trp metabolites detected in mouse sera were assessed for their ability to activate the mouse AHR using a stably transfected luciferase reporter construct in the Hepa1.1 cell line. Nine Trp metabolites and the Trp metabolite Pool were used to treat Hepa1.1 cells for 4 h. Metabolite activity was compared to VEH treatment. Metabolite concentrations were based on serum samples collected from C57BL/6 mice fed a standard laboratory chow diet shown in Table 2. Statistical significance was determined using one-way ANOVA. The data are mean ± SEM. *: p-value < 0.05; **: p-value < 0.005; ***: p-value < 0.0005; ****: p-value < 0.0001. ns: not significant.

**Table 3.** Comparison of serum Trp metabolite concentrations in conventional mice fed semi-purified and standard chow diets.

| Metabolite | Mean abundance ± S.D. (μM) | |
| --- | --- | --- |
|  | Semi-purified diet | Standard chow diet |
| **5-hydroxyindole-3-acetic acid** | 0.52 ± 0.13 | 0.66 ± 0.15 |
| **Indole-3-acetic acid** | 0.42 ± 0.05<sup>a</sup> | 0.68 ± 0.21 |
| **Indole-3-acrylic acid** | 0.10 ± 0.02<sup>a</sup> | 0.31 ± 0.10 |
| **Indole-3-carboxaldehyde** | 0.19 ± 0.07 | 0.19 ± 0.02 |
| **Indole-3-lactic acid** | 1.12 ± 0.18 | 1.66 ± 0.53 |
| **Indole-3-propionic acid** | 0.12 ± 0.10<sup>a</sup> | 4.71 ± 1.48 |
| **Kynurenic acid** | 0.19 ± 0.07 | 0.24 ± 0.09 |
| **Kynurenine** | 1.63 ± 0.22 | 1.10 ± 0.28 |
| **2-Oxindole** | 0.06 ± 0.08 | 0.09 ± 0.05 |

### Tryptophan metabolite profiling of human sera

Trp metabolites circulating in serum could provide insight into homeostatic activity of AHR, and we wished to know if the observations in mice could be replicated in humans. Previous work by our group identified AHR activators in human feces, yet it is not clear if these activators are also present in circulation. To address this question, we quantified Trp metabolites in human serum. To mitigate the variable effect of diet on metabolite profiles, we analyzed sera samples collected from study participants fed a defined diet.^25^ As with the previously discussed mouse sera, targeted metabolomics was performed on serum samples using LC-MS and GC-MS (Table S2). Six Trp metabolites, IPA, IAA, ILA, I3C, KYN, and KA, were detected in human serum at considerably varied concentrations (Fig. 4). Additionally, many metabolites were highly variable across the cohort of trial participants (Table 4). IPA was the second most variable metabolite (CoV: 0.571) and had the greatest range of the metabolites with concentrations detected between 0.443-8.417 μM. KA was the most variable (CoV: 0.763), but least abundant metabolite. The abundances of KA and KYN are positively correlated (Fig. S2, Table S4) which may be explained by a necessary coupling due to the kynurenine pathway, while positive correlations between I3C and all other detected metabolites, except IPA, is unclear. We calculated the sum of all six Trp metabolites across study participants to provide broader insights into individual variability. The wide range of Trp metabolite concentrations (5.52-13.32 μM) added together further illustrates the vast individual variation in Trp metabolite profiles, which emphasizes the need for a substantial sample size to establish a pattern.

**Figure 4.**
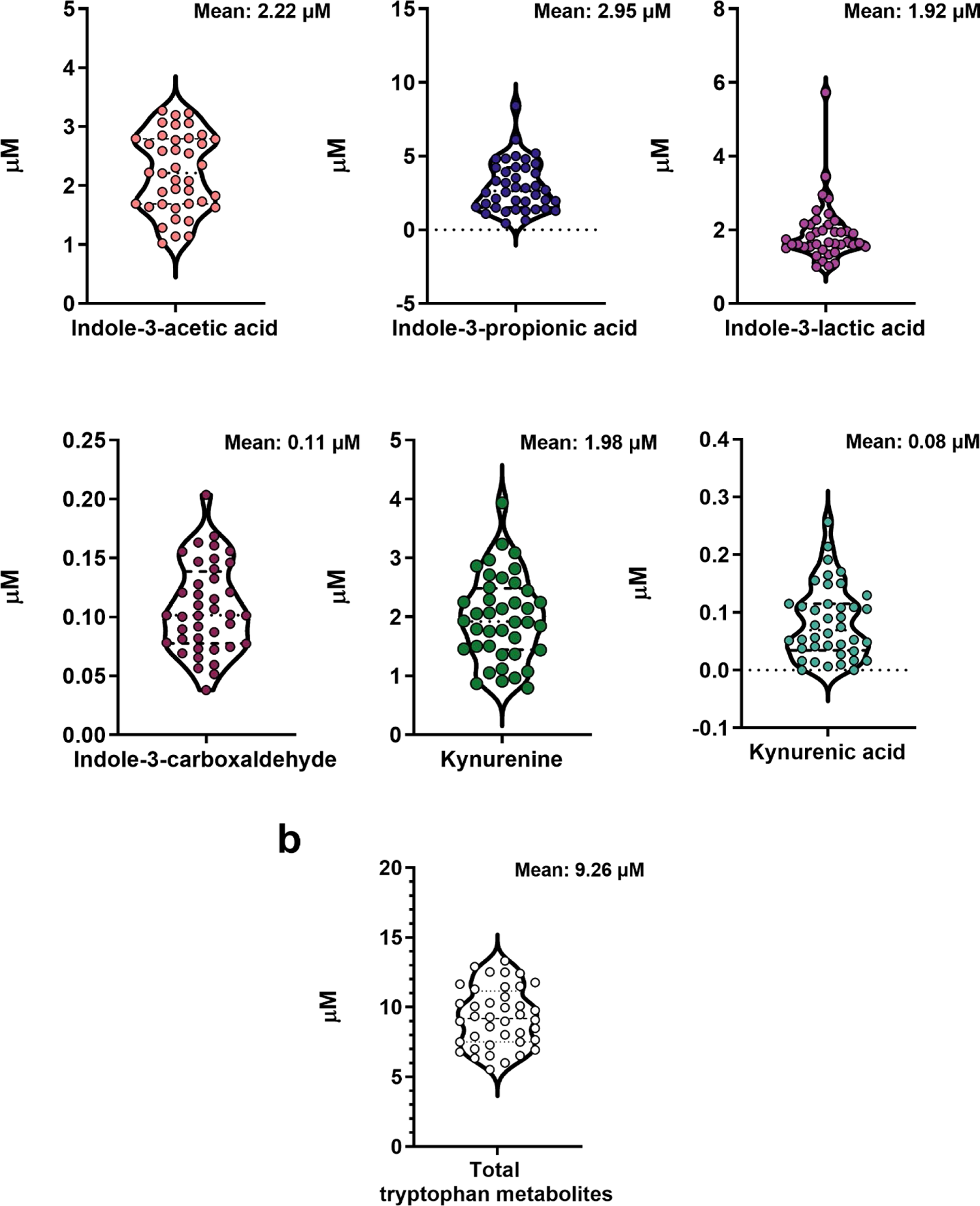
Concentrations of six tryptophan metabolites in human circulating blood serum. LC-MS analysis was used to quantitate tryptophan metabolites (a) Concentrations of six individual Trp metabolites were identified in circulating blood serum collected from study participants provided a controlled diet. (b) Total concentration of Trp metabolites. Each violin plot displays the median value (---) and Q1 and Q3 (···). n = 40.

**Table 4.** Descriptive statistics of human serum metabolites

| Metabolite | Mean ± S.D.<br>(μM) | Range (μM) | Coefficient of<br>Variation | Skewness | Std. Error of<br>Skewness |
| --- | --- | --- | --- | --- | --- |
| Indole-3-propionic acid | 2.954 ± 1.686 | 0.443-8.417 | 0.571 | 0.979 | 0.374 |
| Indole-3-acetic acid | 2.222 ± 0.655 | 1.025-3.274 | 0.295 | -0.09 | 0.374 |
| Kynurenine | 1.981 ± 0.727 | 0.794-3.934 | 0.367 | 0.4 | 0.374 |
| Indole-3-lactic acid | 1.917 ± 0.808 | 0.998-5.731 | 0.421 | 2.963 | 0.374 |
| Indole-3-carboxaldehyde | 0.107 ± 0.038 | 0.038-0.204 | 0.35 | 0.43 | 0.374 |
| Kynurenic acid | 0.083 ± 0.063 | 0-0.257 | 0.763 | 0.782 | 0.374 |

### Circulating tryptophan metabolites generate human AHR activity

Given species differences in the AHR ligand binding pocket, metabolism, and the composition of the gut microbiome, it was unclear if the previously discussed mouse AHR-based activity assays would produce similar results in a human model. To address this question, we used human cell culture models to examine AHR activation potential of the Trp metabolites detected in human sera. We evaluated the potential of the six detected metabolites to activate the AHR using the mean concentrations of each metabolite (IPA: 2.95 μM; IAA: 2.22 μM; ILA: 1.92 μM; I3C: 0.11 μM; KYN: 1.98 μM; KA: 0.08 μM) to assess potential physiologic activation. We also generated a mixture comprised of the mean concentrations of all six detected metabolites, referred to as the “Pool” (Fig. 5a), to provide a more biologically relevant model for assessing systemic AHR activity. AHR activity was measured using a human AHR-dependent luciferase reporter assay system (HepG2 40/6) following 4 h exposure to metabolite treatments (Fig. 5b). The Pool, KA, KYN, and IAA generated significant activity relative to the vehicle treatment, with a 2 or 3-fold induction. Interestingly, the sum of individual metabolites generates far more activity than is seen when cells are exposed to the metabolite Pool, suggesting an antagonistic effect. Furthermore, while the removal of a single metabolite from the Pool does not significantly alter activity (Fig. 5c) relative to the complete Pool; the removal of two dominant metabolites, KA and KYN, does significantly diminish activity. Strikingly, the presence of IAA in this treatment does not protect against a loss of activity. This further supports an antagonistic relationship between the six Trp metabolites and AHR activity. However, it should be noted that the six Trp metabolites were determined to not have cytotoxic effects (Fig. S3).

**Figure 5.**
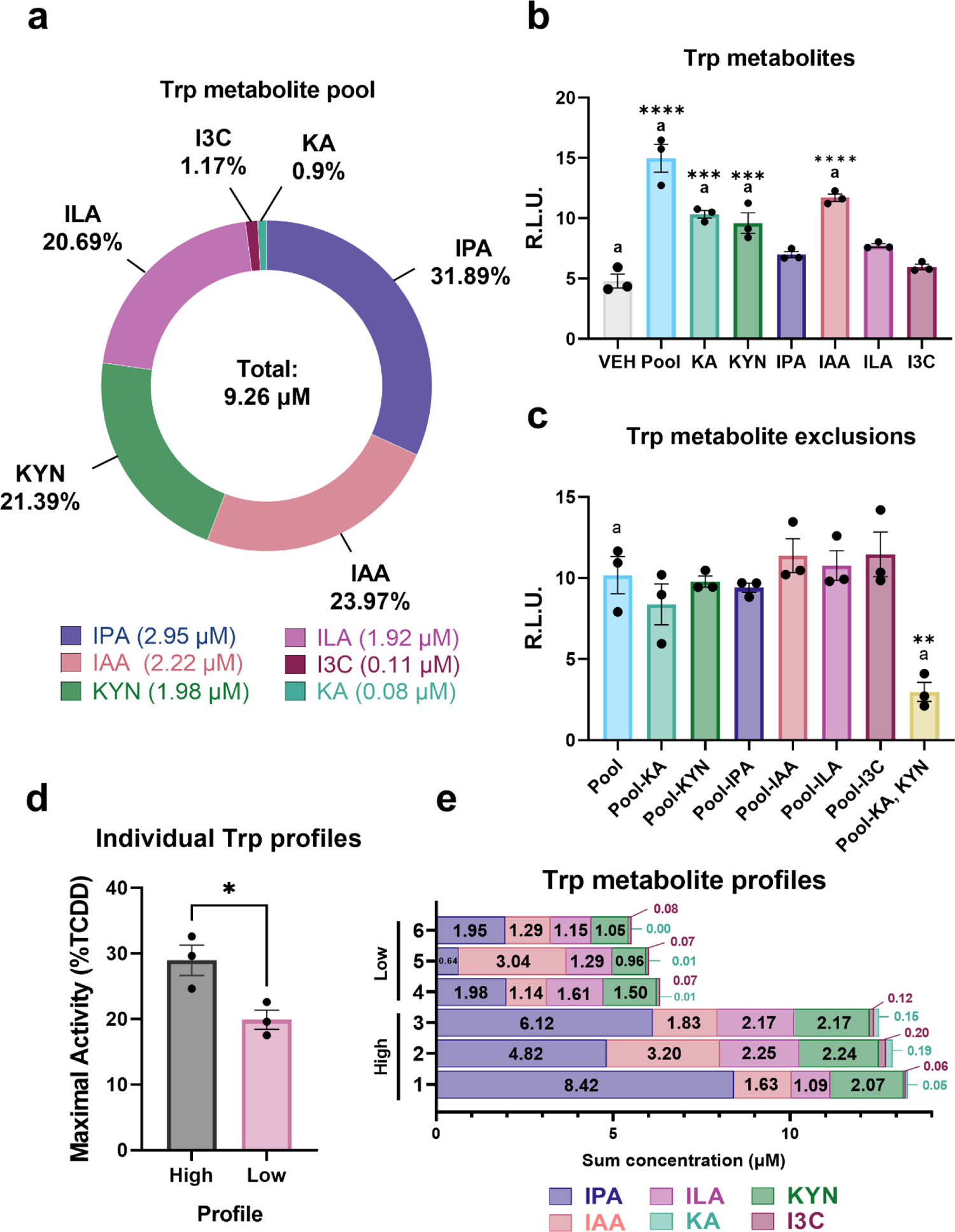
Circulating tryptophan metabolites generate AHR activity at physiological concentrations. Trp metabolites and Pools were assessed for their ability to activate the human AHR using a stably transfected luciferase reporter construct or *CYP1A1* induction in the HepG2 40/6 cell line. (a) The composition of the serum metabolite “Pool” determined by the mean of each tryptophan metabolite. (b) Six tryptophan metabolites at the mean concentration detected in serum and the Pool were used to treat HepG2 40/6 cells for 4 h. Metabolite-generated activity was compared to VEH control. (c) Specific metabolites were excluded from the Pool and used to treat HepG2 40/6 cells for 4 h. Activity from VEH control was subtracted from experimental treatments and AHR activity was compared to the complete Pool treatment. AHR activity was determined using a luciferase reporter system. (d) HepG2 40/6 cells were treated for 4 h with Trp metabolite profiles and AHR activity was measured by the relative expression of *CYP1A1* to *ACTB*. Data are normalized to a saturating dose of TCDD (10 nM) and VEH (0.1% DMSO, final concentration) treatment, and presented as a percentage of maximal AHR activity. (e) Metabolite concentrations representing the Trp metabolite profiles from six individual trial participants, grouped into “High” and “Low” groups based on total metabolite abundance. Statistical significance was determined using one-way ANOVA or unpaired *t-*test. The data are mean ± SEM. *: p-value < 0.05; **: p-value < 0.005; ***: p-value < 0.0005; ****: p-value < 0.0001.

Guided by the potential for an agonist/antagonist dynamic observed in the Trp mixture, we predicted that the total concentration of Trp metabolites would affect the degree of AHR activity. To test this hypothesis, we replicated the Trp metabolite profile of individual trial participants with the three greatest (**1.** 13.32 μM, **2.** 12.9 μM, **3.** 12.53 μM) and lowest total concentrations (**4**. 6.34 μM, **5.** 6.01 μM, **6.** 5.52 μM) of examined metabolites to assess induction of the prototypical AHR target gene, *CYP1A1* (Fig. 5d, e). The three profiles with the highest and lowest total metabolite abundances were grouped accordingly and induced a statistically significant difference in AHR activity. Considering the substantial variation in total metabolite abundance, the surprisingly small difference in AHR activation may suggest that multiple factors may affect AHR activation, but metabolite affinity for the AHR and potential competitive binding dynamics may be a leading factor. We sought to further investigate this question by examining the affinities of AHR for each of the six Trp metabolites.

### Tryptophan metabolites directly interact with the AHR

We assessed the affinity of the six Trp metabolites for the AHR by exposing HepG2 40/6 cells to a range of concentrations for each metabolite (Fig. 6). While the luciferase reporter system is not necessarily reflective of a realistic physiological response, it does allow for relative comparisons of activation potential. EC_25_ values demonstrate that KA is the most potent of the six metabolites with an EC_25_ value of 40.8 nM. The EC_25_ values of KYN (1.76 μM) and IAA (2.92 μM) rank these metabolites the second and third most potent metabolites, respectively. Further, the EC_25_ values for KA, KYN and IAA fit within the range of concentrations that were detected in human sera, suggesting that these values are physiologically possible. Meanwhile, the EC_25_ values for IPA (19.2 μM), ILA (25.9 μM), and I3C (109.8 μM) are far above the range of concentrations that were detected in human sera. These data imply that these metabolites are unlikely to directly generate homeostatic AHR activity at typical physiological concentrations.

**Figure 6.**
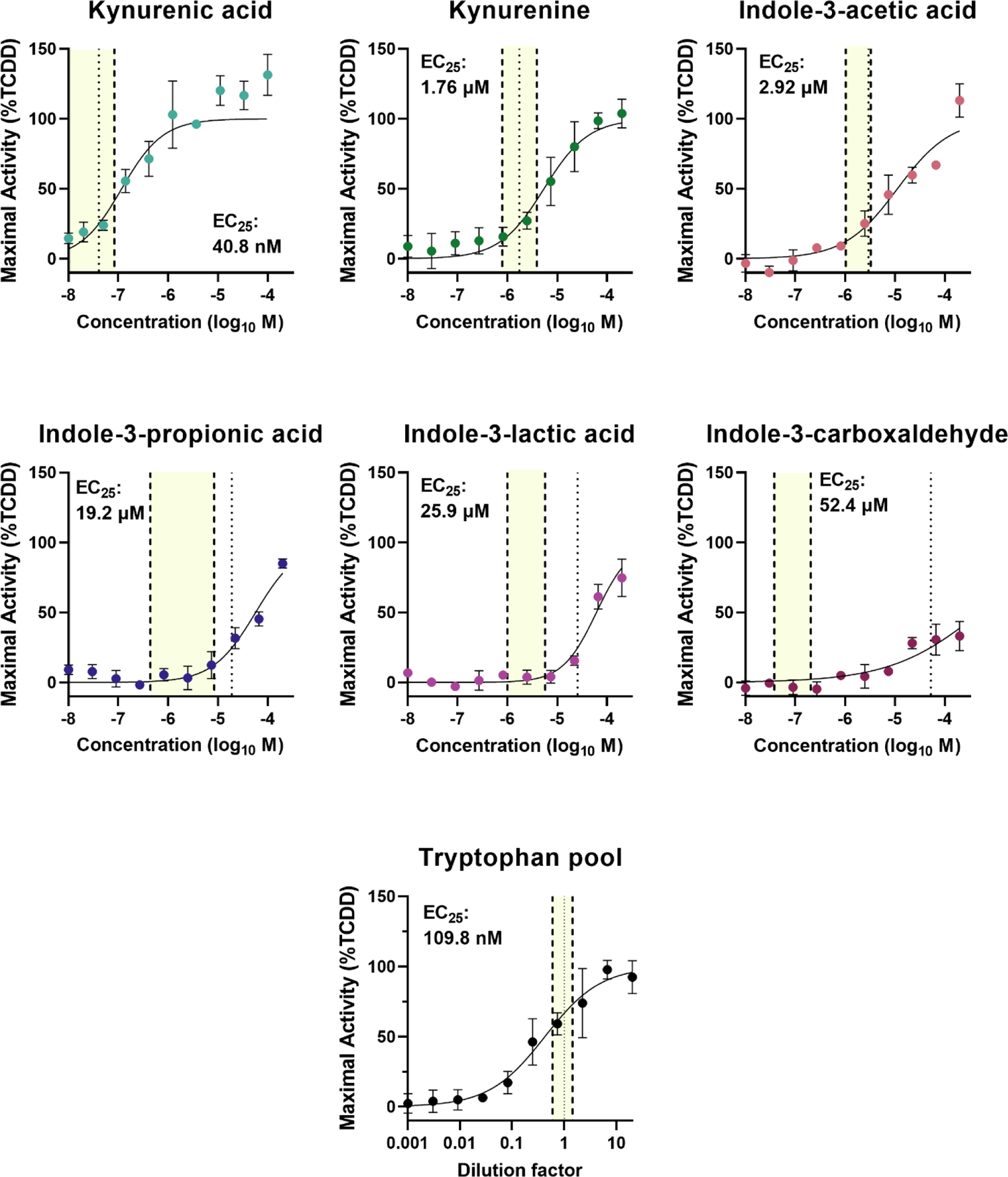
AHR activity in response to multiple treatment doses of tryptophan metabolites. Trp metabolites were assessed for the ability to induce AHR activity across a range of treatment concentrations using a stably transfected luciferase reporter construct in the HepG2 40/6 cell line. AHR activity was determined using a luciferase reporter system following a 4 h treatment time. Data are normalized to a saturating dose of TCDD (10 nM) and VEH (0.1% DMSO, final concentration) treatment, and presented as a percentage of maximal AHR activity. EC_25_ values were generated by nonlinear regression and represented in the figure by a dotted line (···). The range of quantified Trp metabolites from analyzed human serum are represented by dashed lines (- - -), and the range is depicted by the yellow fill.

Although we observed AHR activity in response to treatments with the examined metabolites, it was unclear if these compounds are truly AHR ligands which directly interact with the ligand binding pocket of the receptor. We sought to address this question with a cell-based AHR ligand competition binding assay.^26^ This is a modification of the gold standard of AHR ligand classification that relies on displacement of a high affinity radioactive ligand from the ligand binding site, but incorporates the complexity of cellular biology, including metabolite uptake, efflux, and protein binding. This assay, however, is limited by the potential of further metabolism of the Trp metabolites prior to binding with the AHR. An assay length of 30 min was chosen to mitigate further metabolism. The result of the assay showed that all six metabolites can compete with the photoaffinity ligand (PAL) at 10 μM concentrations (Fig. 7). Importantly, the Pool also demonstrated that it competed with the PAL for binding to the AHR ligand binding site. Furthermore, ligand docking simulation using Autodock Vina provides additional evidence that the Trp metabolites are indeed interacting with the ligand binding site of AHR (see Methods, and Fig. S4). Computational docking analysis of the six Trp metabolites (Fig. 8) revealed that each compound can bind the PAS B domain of the AHR with low micromolar to high nanomolar affinity in the same primary ligand binding pocket, defined by indirubin.^27^ ILA (0.81 nM) and KA (0.95 nM) demonstrated the highest computational affinity for the AHR (Table S5), but this did not correlate completely with AHR ligand binding assay data obtained in HepG2 40/6 cells (Fig. 5). However, all six Trp metabolites bound the AHR in the same hydrophobic groove as indirubin (see Fig. S5), with several forming unique electrostatic interactions with Fα helical residue Ser-336, which was not a well-defined element of the primary indirubin binding pocket.^27^ The formation of this unique contact with helix Fα stabilized the recognition of the smaller, Trp-related ligands, even in the absence of hydrophobic contacts with Leu-353, the important Gβ-sheet residue that appears to define the opening of the primary ligand binding pocket. Although data from our cell-based assays would imply differences in binding affinity, simulated binding energies and dissociation constants are similar across the six metabolites. This is not surprising given the structural similarities between these compounds. Nevertheless, the combination of modeling and biological activity data clearly supports the concept that the Trp metabolites are direct AHR ligands.

**Figure 7.**
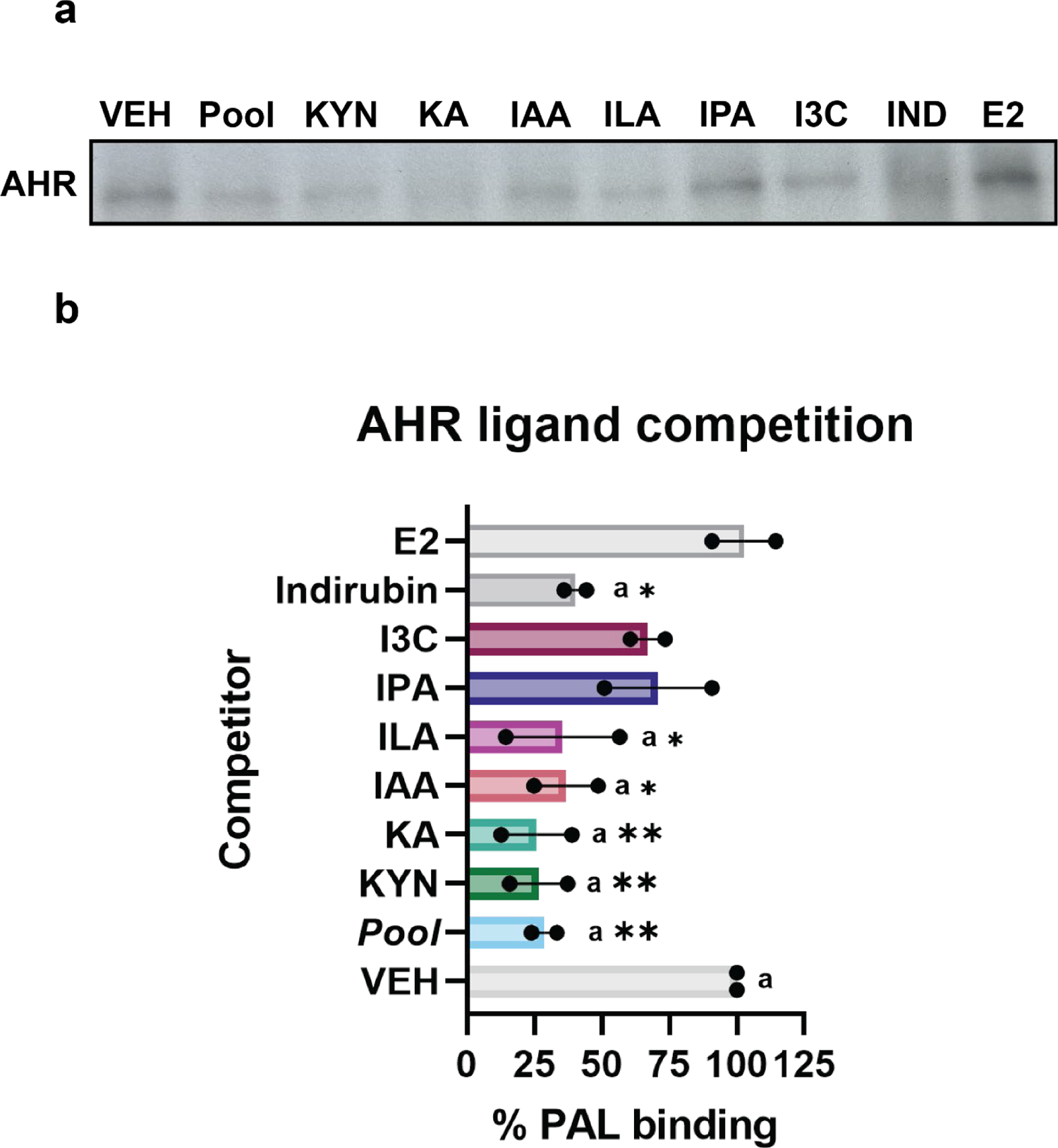
Serum tryptophan metabolites prevent binding of radioactive photoaffinity ligand. Trp metabolites were assessed as AHR ligands by binding to the ligand binding pocket of AHR and preventing the binding of the high affinity photoaffinity ligand 2-azido-3-[^125^I]iodo-7,8-dibromodibenzo-*p*-dioxin (PAL). HN30 cells were with treated 10 μM Trp metabolite and the metabolite Pool (9.26 μM, total Trp metabolite concentration; italicized) prior to the addition of 2 pmoles PAL. Cell lysates were processed and subjected to tricine SDS-PAGE, transferred to PVDF membranes, and AHR band intensity was quantified using densitometry. Kynurenic acid (KA) is a known AHR ligands. Samples were normalized to AHR abundance. (a) PAL competition assay with the Trp Pool and individual metabolites metabolites; all reduced radioligand binding. (b) Quantitative graphic of the results from panel a. The data are mean ± SEM. *: p-value < 0.05; **: p-value < 0.005.

**Figure 8.**
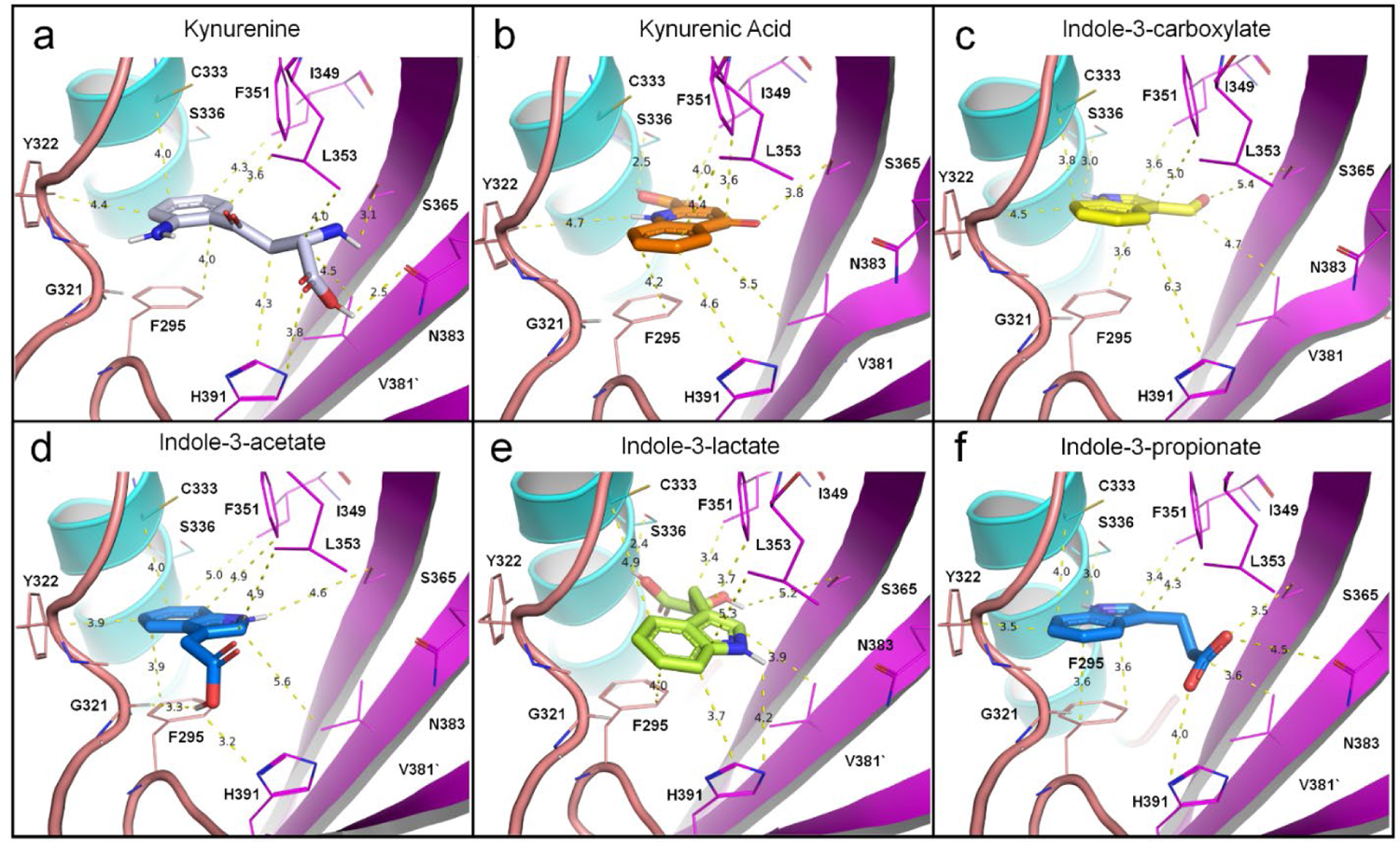
Trp metabolite docking in the cryo-EM model of the human AHR PAS B domain. The substrate binding properties of kynurenine, kynurenic acid and four tryptophan-related metabolites (Indole-3-acetate, Indole-3-carboxylate, Indole-3-lactate, Indole-3-propionate) were studied using the cryo-EM structure of the AhR (7ZUB) and Autodock Vina (see Supplementary Table 1; see Methods). (a) Kynurenine (KYN) docks deeply into the primary, ligand binding domain of the AHR with low micromolar affinity (−7.5 kcal/mol; KD = 3.07 μM; grey stick). KYN engages the primary, indirubin binding site in close contact to key active site residues: F295, Y322, C333, I349, F351, L353, S365, V381, N383 and H391. Electrostatic and hydrophobic bond distances are shown (yellow dashes). (b) Kynurenic Acid (KA) displayed over 3-fold higher affinity to the PAS B domain as KYN (−8.2 kcal/mol; KD = 0.95 μM; orange stick). KA docking revealed balanced interactions among the ligand and both sides of the closed, active site, including a strong electrostatic contact with Fα helix residue S336. (c) Docking results for Indole-3-carboxylate (I3C) revealed over 4-fold lower affinity for the AHR compared to KA (−7.3 kcal/mol, KD = 4.29 μM; yellow stick). I3C preferentially docked in the distal pocket as well, but showed reduced interactions with L353, N383 and H391. (d) Indole-3-acetate (IAA) displayed nearly 2-fold higher affinity for the AHR than I3C (−7.6 kcal/mol, KD = 2.60 μM; dark blue stick), docking the left side of the distal binding pocket via strong interactions with key residues G321, C333, Y322 and H391. (e) Indole-3-lactate (ILA) docked the AHR with high nanomolar affinity (−8.3 kcal/mol; KD = 0.81 μM; light green stick), displaying the highest affinity of all 6 Trp-related ligands tested. ILA binds deeply into the primary pocket, forming close contacts with all key active site residues shown, except for G321, Y322 and N383. (f) Indole-3-propionate (IPA) bound the AHR with similar affinity as ILA and KA (−8.1 kcal/mol; KD = 1.13 μM; purple stick), but deeper in the distal pocket, forming strong contacts with F295 and S336, but not L353, which may be a strong determinant of proper ligand recognition in the AHR, as needed for efficient receptor transactivation.

**Figure 9.**
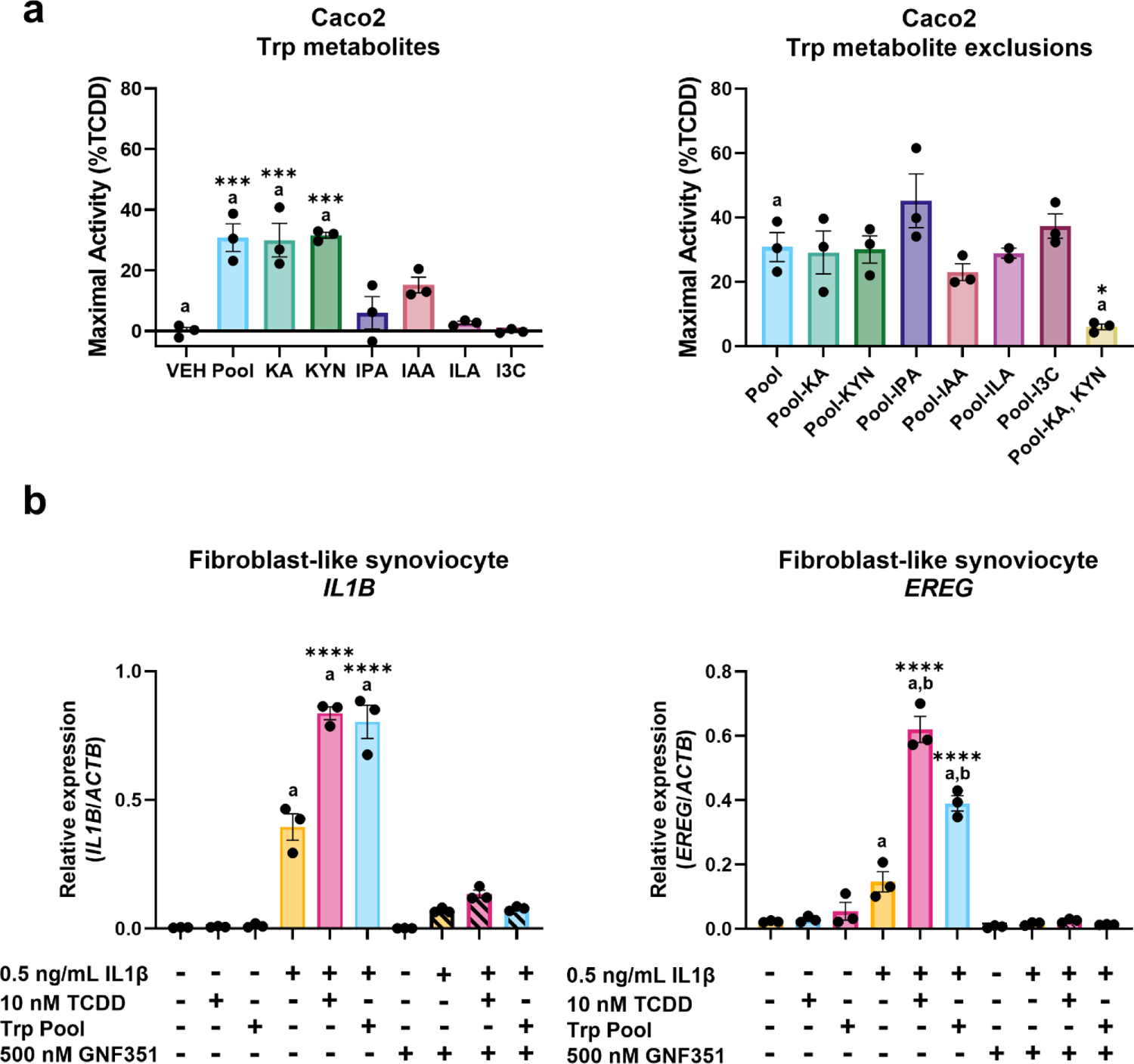
Circulating tryptophan metabolites influence AHR-mediated gene expression in human disease models. (a) AHR target gene, *CYP1A1*, induction in response to 4 h treatment with Trp metabolites in human colorectal adenocarcinoma cells (Caco2) compared to VEH or Pool treatment. Data presented as a percentage of maximal AHR activity (10 nM TCDD). (b) Primary human fibroblast-like synoviocytes isolated from heathy patients were co-treated for 24 h with IL-1β and the Trp pool or pretreated with AHR antagonist (GNF351) prior to treatments. Induction of inflammatory mediators, *IL1B* and *EREG*, was measured. Gene induction was determined by real-time qPCR. Data are normalized to a saturating dose of TCDD (10 nM) and VEH control, and presented as relative to *ACTB* expression. Statistical significance was determined using one-way ANOVA. The data are mean ± SEM.

### Trp metabolites activate AHR in disease models

We wanted to assess the influence of detected Trp metabolites on models for diseases that have been linked to AHR to provide insight into the physiological relevance of the Trp Pool metabolites. First, we used Caco2 cells to model the impact of Trp metabolites on a cancer model. Previous studies have primarily investigated the role of individual metabolites in isolation, yet the data presented here indicate that many AHR activators likely circulate throughout the body together. Caco2 cells were exposed to individual metabolites at mean serum concentrations, the complete Trp metabolite Pool, and variations of the Pool (excluding individual metabolites) for 4 h prior to RNA isolation. Quantification of AHR target gene, *CYP1A1*, transcripts by qPCR established that only KA, KYN and the Pool generate AHR activity significantly different from the vehicle treatment. Moreover, removing a single metabolite from the Pool does not alter AHR activity, yet removing the two most potent AHR activators, KA and KYN, from the Pool substantially decreases AHR activity.

Second, we investigated the impact of the Trp metabolite Pool by exposing a primary cell line of fibroblast-like synoviocytes (FLS) collected from healthy patients to the Trp metabolite Pool. FLS are used as a model in the study of rheumatoid arthritis, an AHR-linked disease, and we have previously shown a synergistic effect on the expression of inflammatory mediators when cells are exposed to IL1β and AHR activators.^4^ FLS were co-treated with 0.5 ng/ml IL1β and a saturating dose of TCDD (10 nM) or the Trp Pool for 24 h prior to RNA isolation, to quantify expression of inflammatory mediator genes, *IL1B* and *EREG*. Additionally, FLS were pre-treated with 500 nM GNF351, an AHR antagonist, to demonstrate the necessity of the AHR in synergistic expression of these genes. The IL1β and Trp Pool co-treatment generated a comparable 185-fold induction in *IL1B* expression to the 192-fold increased produced by the IL1β and TCDD. Expression of *EREG* significantly increased with both co-treatments compared to IL1β alone, however, the IL1β and TCDD co-treatment produced a greater 27-fold induction in *EREG* compared to a 17-fold change generated by IL1β and Trp Pool.

## Discussion

Previous work from our group and others have quantified the abundance of potential AHR-activating Trp metabolites from feces and cecal contents, however, it is not clear whether these represent concentrations to which the intestinal epithelial lining is exposed or circulate in the bloodstream. The extraction of water from waste material in the colon specifically concentrates hydrophobic metabolites, therefore, the concentration of many nonpolar metabolites will likely be higher in this context than the concentrations that intestinal epithelial cell are exposed to. Further, the exposure of intestinal epithelial cells to metabolites will not be comparable to the cell exposure to these metabolites throughout the body. Transport, metabolism, and compound clearance represent possible mechanisms that would reduce circulating metabolite concentrations. To address these concerns, we chose to quantitatively profile circulating metabolites to provide the foundation for investigating systemic or basal AHR activity. We then focused on quantifying Trp metabolites that we previously demonstrated are AHR activators to provide a measurable metabolite profile.^20^ The promiscuity of the AHR ligand binding pocket means that hitherto uncharacterized activators will inevitably be discovered and further complicate the dynamics presented here. Nonetheless, the data provide a basis for more complex models that are necessary to understand the nuances of physiological AHR activation.

To investigate systemic AHR activity, we used mouse models to test specific hypotheses, however, we chose to minimize our reliance on mouse models because they do not accurately portray the human AHR mediated activation by Trp metabolites. Similar to the human AHR, mouse AHR activity is significantly induced in response to the mouse Trp Pool however, unlike human AHR, it is primarily responsive to a single metabolite, KYN. This suggests the known evolutionary changes in the human AHR binding pocket allow for greater recognition of KA and indole-containing compounds.^28,29^ Furthermore, the greater relative affinity of the endogenously produced KA for human AHR, may represent an evolutionary change allowing for greater control over AHR activity. The differences in receptor affinity highlight the importance of appreciating the structural differences between mouse and human AHR and limit the use of mouse models to explore this topic. Beyond AHR structural differences, the mouse microbiome is markedly different from human and this also influences the products of metabolism.^30,31^ Moreover, our lab has previously shown that, despite high expression of AHR in the colon of conventional C57BL/6J mice and high concentrations of Trp metabolites, there is little to no *Cyp1a1* expression.^32^ Taken together, these observations imply that mouse AHR is relatively insensitive to changes in Trp metabolites in the context of detectable serum concentrations. These distinctions present the biological challenges limiting the accuracy of mouse models to provide specific insights into physiologically induced human AHR activity.

Demonstrating AHR activation in response to a compound is often thought to be sufficient evidence to assume that a specified compound is an AHR ligand. However, some compounds, such as serotonin and butyrate, generate activity without directly interacting with the AHR.^33^ The gold standard for identifying an AHR ligand is a radioligand competition binding assay to demonstrate compound interaction with the AHR binding pocket. We employed a cell-based radioligand competition assay to evaluate all six metabolites for their ability to compete with the high affinity PAL. Compared to an *in vitro* radioactive ligand competition assay, the cell-based method inherently considers transport, efflux, and other cellular functions that may interfere with a compound from binding to the receptor, providing a biologically relevant context. However, this assay is limited in that it allows further metabolism of treatment compounds. We sought to address this limitation with both a short treatment time and with supporting evidence from additional methods. We generated dose response curves for the six Trp metabolites to characterize AHR responsiveness to each metabolite. Lastly, we used ligand binding simulations to model the binding interaction of each metabolite with the AHR ligand binding pocket. Simulated ligand docking similarly posed all six Trp metabolites and in the same plane as the well-known AHR ligand, indirubin, supporting that the six Trp metabolites are indeed AHR ligands. Despite similar binding energies and dissociation constants, suggesting little theoretical difference in activation potential between the six metabolites, this is not reflected in cell-based activity assays. This discrepancy underscores the importance of studying potential AHR ligands in a biological context that considers metabolism and other cellular functions, because theoretical binding potential does not guarantee actual receptor activation. Nonetheless, docking simulation analysis upholds the hypothesis that these six Trp metabolites can act as AHR ligands. Support from these three methods provide evidence that the examined metabolites are directly interacting with the AHR and are, indeed, genuine AHR ligands.

Research into Trp metabolism across taxonomic kingdoms has created confusion about the source of specific metabolites, especially indole-containing compounds. It is often assumed that the production of a Trp metabolite at relatively high concentrations by gut bacteria means that bacteria are the only or primary source for the host. Our data suggest this is not always the case. IAA is often depicted as a product of microbial metabolism, despite long recognition as an essential plant growth hormone. Further, the data presented here from GF mice indicate that IAA is produced independent of bacteria and is supported by previous publications reporting human enzymes that can produce IAA.^34^ This confusion is seen in previous reports of ILA or I3C as well, while our samples collected from GF mice indicate that these compounds circulate in a host without the presence of bacteria.^35,36^ Yet, we are confident that 2-oxindole, indole-3-acrylic acid, and IPA are microbial products and represent metabolic products of the microbiome that can circulate in a host. Not only are these below the limit of detection in GF mice, but the published literature supports microbial production of these compounds.^14,37,38^ Notably, the presence of indole-3-acrylic acid and IPA are likely partially coupled because indole-3-acrylic acid is a known intermediate in a microbial metabolic pathway that terminates in IPA.^14^ Given that IPA circulates in humans (HMDB ID: HMDB002302), it is possible that microbial metabolism may contribute to the circulating pool of IAA, ILA, or I3C, but further work is needed to understand this dynamic.^39^ Strikingly, metabolites found in high concentration in feces, e.g. indole, were undetectable in serum.^20^ This disparity highlights the value of direct metabolite analysis of tissues or biofluids in addition to proxy measurements, like feces, when measuring the concentrations of AHR ligands.

Study of AHR-activating Trp metabolites often leads to suggestions that changing the composition of the gut microbiome may alter AHR activity, but this proposition may be more challenging than it appears at first glance. Bacterial colonization of the gastrointestinal tract depends on multiple competitive dynamics, thus, producing an environment that is difficult to control.^40,41^ In contrast, designing an individual’s diet to achieve a clinical outcome is long standing practice. We wanted to demonstrate the possibility that diet affecting Trp metabolism could alter potential clinical outcomes. We observed that placing mice on a semi-purified diet does change Trp metabolite abundances. The two diets produced differences in serum concentration of three of the nine metabolites, two of which are microbially produced indole-3-acrylic acid and IPA. This suggests that the semi-purified diet can suppress specific microbial metabolites, and these compounds appear particularly sensitive to change. The fact that 2-oxindole concentration showed no change suggests that altering diet may only affect specific microbial pathways. Similarly, the significant reduction in IAA in mice fed a semi-purified diet indicates that host Trp metabolism may be altered by diet, but changes in microbial production of IAA remains a possibility as well. Although the removal of phytochemicals from the semi-purified diet may offer a possible explanation, the cause of these metabolic changes is unclear and requires additional research. Whereas the abundance of serum Trp metabolites can be modulated by non-pharmacological means, the influence this would have on AHR activity is difficult to predict at this point.

We used a human AHR-dependent reporter system (HepG2 40/6) to specifically assess the potential for each metabolite to activate the AHR at the identified mean concentration. This system is limited in its capacity to dissect biological responses but is adequate for relative comparisons and provides broad insights into how these metabolites activate AHR. This assay indicates some indoles, IPA, I3C, and ILA, are not significantly contributing to AHR activation potential at physiological concentrations. Measuring *CYP1A1* expression induced by the Trp Pool supports the notion that not all Trp metabolites generate activity at homeostatic concentrations. Notably, KA is a downstream product of KYN metabolism, coupling these two important AHR agonists.^42^ However, this may be dependent on the observed cell type, given that the expression profile of kynurenine pathway enzymes will differ between cell types and physiological conditions. Determining the role of this pathway in producing AHR ligands may be critical to understanding systemic AHR activity and warrants further investigation. The significant positive correlation between KA and KYN presents another challenge of disentangling individual metabolite activity due to possible continued metabolism within cells. Further, relative to the commonly studied industrial pollutants (e.g., polychlorinated biphenyls) or dietary compounds (e.g., indole-3-carbinol) that act as AHR ligands, these six Trp metabolites are low affinity ligands, and, as such, may serve as a regulatory mechanism. Saturation of the AHR by Trp metabolites could prevent or hinder AHR-mediated response to environmental toxins or immune challenge. Therefore, the low affinity of these metabolites to the AHR may maintain basal homeostatic AHR activity while permitting a strong response to environmental challenges.

The role of AHR in cancer is well documented, so we investigated whether the six identified metabolites would generate AHR activity in a colorectal cancer model (Caco2).^43,44,45,46^ Our results indicate that the examined metabolites do activate AHR in Caco2 cells leading to a change in *CYP1A1* expression. In some types of cancers this may provide a protective effect for a tumor by contributing to immune system suppression. Host genetics, metabolic dysregulation, or diet may exacerbate this effect, producing greater AHR activation and potentially influencing clinical outcomes in patients. A comparison of activity produced in HepG2 and Caco2 cells also suggests that there may be cell type specific responses to these metabolites.

To further examine this possibility, we exposed FLS to the Pool of Trp metabolites. Previously, we have shown that AHR activation acts combinatorially with IL1β to induce inflammatory mediators, including *IL1B* and *EREG*, in FLS.^4,47^ Co-treatment with IL1β and the Trp Pool led to increased expression of both biomarkers associated with worsened clinical outcomes. Regular or constant exposure to the examined Trp metabolites may provide insights into understanding the development of rheumatoid arthritis, an important AHR-linked disease.^48,49^ The presence of Trp metabolites that activate the AHR leads to the priming of the IL1B promoter facilitating enhanced transcriptional activity in the presence of a pro-inflammatory response, such as NFkB activation.^4^ This model represents potential future directions for AHR-linked disease research. Indeed, Trp metabolites may exacerbate a disease state, or dysregulation of Trp metabolism may lead to the development of disease.

Comparing the results of treating cells with the Trp Pool and individual metabolites demonstrate the difficulty in interpreting the impact of a mixture of Trp metabolites on AHR activity. Our experiments factor in the many cellular processes (metabolism, transport, and competition for receptor binding) that likely act upon the metabolites without providing insight into how these processes are altering activity. Transporters for KA and KYN have been identified, but the uptake mechanisms of the four indole-containing compounds have been poorly studied.^50,51^ The differential expression of transporter proteins in distinct cell types may result in varied intracellular concentrations that affect AHR activity. Additionally, the half-life of these metabolites is likely determined by further metabolism, host clearance and compound stability. Furthermore, the rate of production of these metabolites may be affected by genetics, microbiome composition, and environmental exposures that are unique to each host. This is further complicated by the possibility that metabolite concentrations vary throughout the day as meals are consumed and Trp availability changes.^52,53^ These processes complicate interpretation and, to further understand homeostatic AHR activity there is a need to develop more accurate models. Additionally, the promiscuity of the AHR binding pocket and the diversity of metabolic compounds imply the inevitability of identifying additional AHR activators. Novel activators should not be assumed to have physiological relevance when assessed in isolation, considering the already significant abundances of endogenous ligands. These studies revealed that circulating Trp metabolites capable of mediating AHR are produced by the host and thus do not originate from the gut microbiome, in contrast to the prevailing dogma in the literature. However, it is possible that homeostasis of circulating concentrations is modulated through increased excretion and absorption of Trp metabolites. Future studies should turn to assessing the important host metabolic pathways that result in the production of Trp metabolites that may regulate basal AHR activity. Finally, it is possible that individual differences in Trp metabolite production capable of mediating AHR activation may play a role in chronic diseases.

## Methods

### Chemicals and reagents

All chemicals, unless otherwise stated, were purchased from Sigma (St. Louis, MO, USA). Indole-3-propionic acid and indole-3-lactic acid were from Alfa Aesar (Heysham, UK). AIN93G was purchased from Dyets, Inc., (Bethlehem, PA). LC/MS grade solvents including, methanol and acetonitrile, were purchased from Fisher Scientific (Hampton, NH, USA). The stock solutions of all reference standards for liquid chromatography were prepared in 10% acetonitrile (v/v) containing 1 μM chloropropamide (internal standard). The mixed standard solutions were gradually diluted with 10% acetonitrile (v/v) containing 1 μM chloropropamide for generating the calibration curves. The AHR photoaffinity ligand 2-azido-3-[^125^I]iodo-7,8-dibromodibenzo-*p*-dioxin (PAL) was synthesized as described.^26^

### Human serum samples

Serum samples were collected as described.^25^ Briefly, 40 selected trial participants were fed a standardized isocaloric diet (representative of an average American macronutrient intake) to achieve weight maintenance for four weeks At the end of the diet period and following a 12 h fast (and avoidance of over-the-counter medication), serum was collected, stored at −80C and subjected to analysis by metabolomics.

### Serum sample collection from conventional and GF mice

C57BL/6J wild type mice were originally purchased from Jackson Laboratories (Bar Harbor, ME). Germ-free (GF) C57BL/6J mouse experiments were conducted at the AAALAC-accredited Animal Resource Center at Montana State University. GF mice were reared in standard, sterile (autoclaved cages containing sterile (autoclaved) bedding inside a hermetically sealed isolator with HEPA-filtered airflow and maintained on sterile (autoclaved) water and sterile (autoclaved) food (LabDiet® 5013, Land O’Lakes) ad libitum. Surveillance of the GF condition was done with standard microbial cultivation and molecular biology techniques^46^. Briefly, liquid ‘bug’ traps comprised of a mixture of food and drinking water were left open to the air inside of isolators and observed daily for signs of microbial growth (i.e., turbidity). Stool samples from mice were monitored prior to and throughout experiments for signs of growth on rich media under anaerobic and regular atmospheric conditions (Mueller–Hinton broth and agar plates). Bulk DNA was also extracted from stool samples (DNeasy PowerSoil Pro DNA isolation kit, Qiagen, Hilden, Germany) to serve as a template for PCR using primers targeting the bacterial 16S rRNA encoding gene. Evidence supporting the GF condition was the combination of a lack of observable microbial growth and 16S amplification by PCR. All GF mouse procedures were approved by Montana State University’s Institutional Animal Care and Use Committee. A subset of mice was switched onto a semi-purified AIN 93G from a chow diet for seven days prior to serum collection. Serum samples from conventional and GF mice were collected and stored at −80°C. All animal protocols were reviewed and approved by the Animal Care and Use Committee of Penn State University.

### Quantification of tryptophan metabolites in mouse and human serum

Tryptophan metabolites were quantified by LC/MS based on established method with some modifications.^20^ Briefly, thawed on ice, 25 μL of serum sample, was mixed with 100 μL extraction solvent of ice-cold methanol containing indole-3-acetic acid-d4 and kynurenic acid-d5. Mixture was vortexed and incubated at −20 °C for 30 min. Following centrifugation at 12,000 × g for 15 min at 4°C, 90 μL supernatant was collected, evaporated to dryness (Thermo Scientific, Waltham, MA) and dissolved in 45 μL of 10% acetonitrile containing 1 μM chlorpropamide. After centrifugation at 12,000 × g for 15 min at 4°C, supernatants were transferred to autosampler vials for LC-MS analysis. Quantitative analysis was performed by reverse phase UHPLC using a Prominence 20 UFLCXR system (Shimadzu, C57Columbia, MD) with a Waters (Milford, MA) BEH C18 column (2.1 × 100 mm, 1.7 μm particle size) maintained at 55 °C and a 20 min aqueous acetonitrile gradient, at a flow rate of 250 μL/min. Solvent A was water with 0.1% formic acid and Solvent B was acetonitrile with 0.1% formic acid. The initial conditions were 97% A and 3 % B, increasing to 45% B at 10 min, 75% B at 12 min where it was held at 75% B until 17.5 min before returning to the initial conditions. The eluate was delivered into a 5600 (QTOF) TripleTOF using a Duospray™ ion source (SCIEX, Framingham, MA). Importantly, data was collected in multiple reaction monitoring (MRM) mode, and the mass accuracy was calibrated by calibration solution purchased from SCIEX (Framingham, MA).

### Cell culture conditions

Caco2 cells were grown in αMEM (Sigma-Aldrich, St. Louis, MO, USA) supplemented with 20% (v/v) FBS (GeminiBio Products, Sacramento, CA, USA) and 1% (v/v) penicillin-streptomycin. Cells were seeded into 6-well plates and grown seven days post-confluence prior to treatments to allow for differentiation to occur. HepG2 40/6 cells were generated as previously described, and Hepa 1.1 cells were a gift from Dr. Michael S. Denison.^54^ HepG2 40/6 and Hepa 1.1 cells were grown in αMEM supplemented with 8% (v/v) FBS and 1% (v/v) penicillin-streptomycin, and seeded into 12-well plates 24 h prior to treatment. Primary human fibroblast-like synoviocytes from healthy patients were purchased from and maintained in Synoviocyte Growth Medium (Cell Application, Inc., San Diego, CA). HN30 cells were obtained and cultured as previously described.^55^

### Cell-based luciferase reporter assay

Prior to treatment, culture media was aspirated from HepG2 40/6 cells, which were thoroughly washed with PBS, then αMEM supplemented with 5 mg/mL BSA, 100 U/mL penicillin, and 100 μg/mL streptomycin was added to each well. Cells were treated with TCDD or tryptophan metabolites in DMSO (0.1% final concentration in cell culture). After 4 h incubation, cells were thoroughly washed with PBS, lyzed using 100 μL of lysis buffer (25 mM Tris-phosphate pH 7.8, 2 mM dithiothreitol, 2 mM 1,2-diaminocyclohexane-N,N,N’,N’-tetraacetic acid, 10% (v/v) glycerol, 1% (v/v) Triton X-100), and stored at −80°C until analysis. Luciferase activity was measured using a TD-20e luminometer (Turner Biosystems Inc.) and luciferase substrate (Promega, Madison, WI, USA), per the manufacturer’s instructions.

### RNA isolation and RT-qPCR

Prior to treatment, culture media was aspirated from HepG2 40/6 and Caco2 cells, which were thoroughly washed with PBS, then αMEM supplemented with 5 mg/mL BSA and 1% (v/v) penicillin/streptomycin was added to each well. HepG2 40/6 and Caco2 cells were exposed to Trp metabolites for 4 h. FLS were treated in Synoviocyte Growth Medium for 24 h. RNA was isolated from cells treated with tryptophan metabolites for quantitative real-time PCR (RT-PCR), performed as previously described.^56^ Primers are listed in supplementary table 3.

### Cell-based AHR ligand competition binding assay

HN30 cells were seeded into 12-well plates at 50,000 cells/well and cultured for 48 h in DMEM/F12 medium (Sigma) with 10% fetal bovine serum (Gemini BioProducts), 25 mM HEPES (pH 7.4), 100 U/mL penicillin, and 100 μg/mL streptomycin. The medium was removed, cells washed with Dulbecco’s phosphate buffered saline (PBS, Sigma). Cells in each well were incubated in 500 μl Hanks balanced salt solution (Gibco), 25 mM HEPES pH 7.4, and 5 mg/ml bovine serum albumin for 30 min. Next, cells were treated with AHR ligands, followed by the addition of 2 pmoles of PAL, cells were placed back in the incubator for 30 min. Medium was removed, 500 μL of PBS was added, and cells were exposed to UV light at >302 nm, at a distance of 8 cm, for 4 min using two 15-W UV lamps (Dazor Mfg. Corp. St. Louis, MO). The PBS was removed and cells lyzed in 100 μL of 25 mM MOPS, 2 mM EDTA, 0.02% sodium azide, 10% glycerol and 1% NP40. Lysates were transferred to microfuge tubes and centrifuged at 18,000 x g for 20 min. Supernatants were subjected to tricine SDS-PAGE, and subsequently transferred to PVDF membrane. The radioactive AHR band was visualized with X-ray film and densitometry analysis was performed with ImageJ v. 1.53t^57^.

### Computational docking analysis

The ligand binding affinity of 6 Trp-metabolites, including kynurenine, kynurenic acid, indole-3-carboylate, indole-3-acetate, indole-3-lactate, and indole-3-propionate, were investigated using Autodock Tools 4 and Autodock Vina, and a cryo-EM structure-based computational model of the human AHR PAS B domain.^27,58,59,60^ Docking models for each of the 6 Trp-related ligands were obtained from the PubChem database and optimized for docking using Autodock Tools.^61^ Autodock Vina was run using standard settings, and Grid box parameters generated in Autogrid for a 30-60 Å3 docking grid centered on the indirubin binding site, as described previously.^62^ Docking results were analyzed using Autodock Tools 4 and The PyMOL Molecular Graphics System, Version 2.52 Schrödinger, LLC.^63^ Active site cavity size was calculated using Caver 3.0.^64^ AHR docking model quality was validated using an Autodock Vina re-docking analysis with the structural coordinates of indirubin. RMSD re-docking error was calculated using the program LigRMSD.^65^

### Statistical analysis

All data were compared using one-way ANOVA in GraphPad Prism v. 9.4.1 (GraphPad Software, San Diego, CA, USA) to determine statistical significance between groups. Values of p <0.05 were considered statistically significant (*: p<0.05; **: p<0.01; ***: p<0.001; ****: p<0.0001).

## Acknowledgements

We thank Marcia H. Perdew for critically reviewing the manuscript.

## List of abbreviations

AHR: Aryl hydrocarbon receptor
CoV: Coefficient of variance
GC-MS: Gas chromatography paired with mass spectrometry
GF: Germ-free
FLS: Fibroblast-like synoviocytes
IAA: Indole-3-acetic acid
IAcA: Indole-3-acrylic acid
I3C: Indole-3-carboxaldehyde
ILA: Indole-3-lactic acid
IPA: Indole-3-propionic acid
LC-MS/MS: Liquid chromatography paired with tandem mass spectrometry
KA: Kynurenic acid
KYN: Kynurenine
PAL: Photoaffinity ligand
SD: Standard deviation
SEM: Standard error of the mean
TCDD: 2,3,7,8-tetrachlorodibenzo-*p*-dioxin
Trp: Tryptophan

---

**Figure S1.**
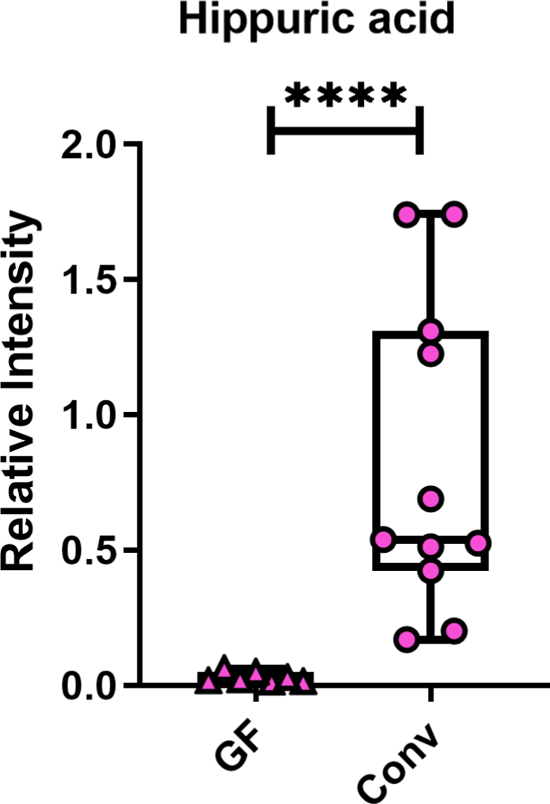
Serum concentrations of hippuric acid were used as a control to demonstrate depletion of bacteria in germ free mice. A statistical comparison was made using a Mann-Whitney test. Each box represents the median value with Q1 and Q3 range, and whiskers describe the maximal and minimal values. Conv, conventional mice; GF, germ-free mice. (****): p-value < 0.0001.

**Figure S2.**
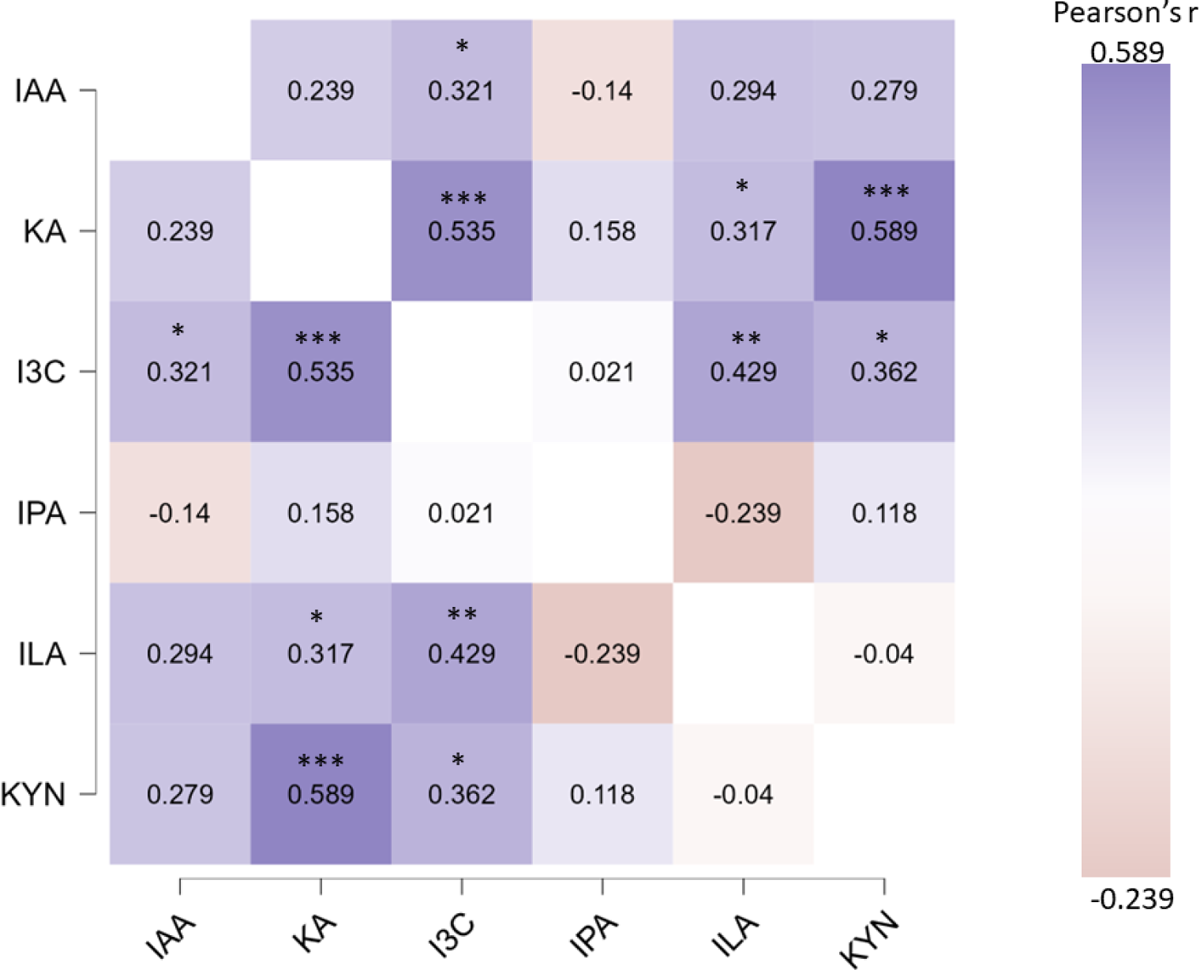
Heatmap of Pearson’s correlations between tryptophan metabolites detected in human serum. Numbers and color indicate Pearson’s r.* p < 0.05; ** p < 0.01; *** p < 0.001.

**Figure S3.**
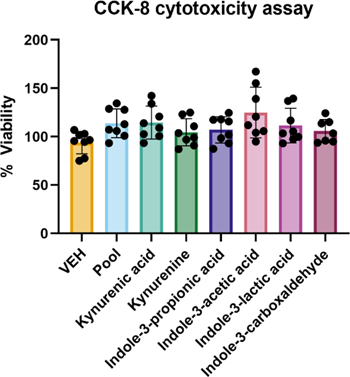
Trp metabolite cytotoxicity assay in Caco2 cells. Cytotoxicity at was assessed using Cell Counting Kit-8 (SIGMA-ALDRICH, Co., St. Louis, Missouri, USA), per the manufacturer’s recommendations, following a 4 h exposure of Caco2 cells to mean serum concentrations of Trp metabolites. Statistical comparisons were determined by one-way ANOVA.

**Figure S4.**
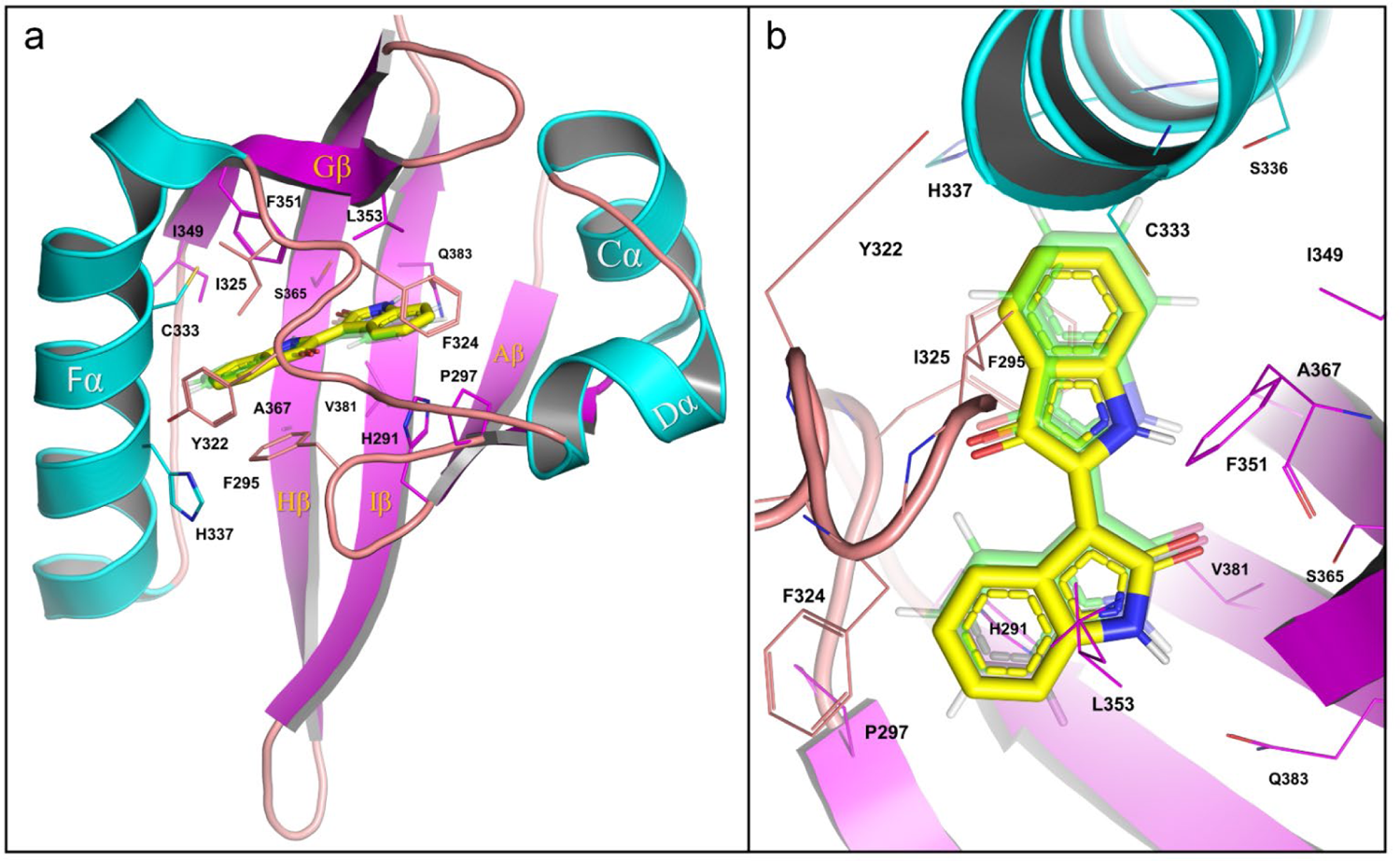
Trp metabolite docking analysis in the cryo-EM structure of the AHR. Ligand binding properties were assessed using Autodock Vina and the PAS B domain of the indirubin-bound Hsp90-XAP2-AHR complex (PDB: 7ZUB [55]; see methods). A.) The PAS B domain of the AHR is shown in cartoon format with α-helical elements C, D and F (in cyan) and β-sheet domains A, G, H, and I (in magenta) highlighted. Key amino acid residues that define the primary ligand binding site are shown (line format), including H291, F295, P297, I325, C333, H337, I349, F351, L353, S365, A367, V381, and Q383. The structural coordinates for indirubin (transparent green stick) are shown superimposed on the low energy, indirubin redocking result (−12.8 kcal/mol; K_D_ = 0.44 nM; see Supplemental Table 5). B.) Autodock Vina redocking results for indirubin in the AHR active site demonstrated high precision in recapitulating ligand recognition (RMSD = 0.498 Å, see Supplemental Table 1), validating the cryo-EM AHR PAS B domain structure as a suitable model for the comparative ligand binding analysis conducted here. While individual ligands may contribute different enthalpic cues to facilitate closure and transactivation of the receptor, in situ, the indirubin-bound active site provides a tangible, closed conformation for exploring structure-activity relationships that are independent of structural artifacts induced via MD-simulation based ligand docking approaches.

**Figure S5.**
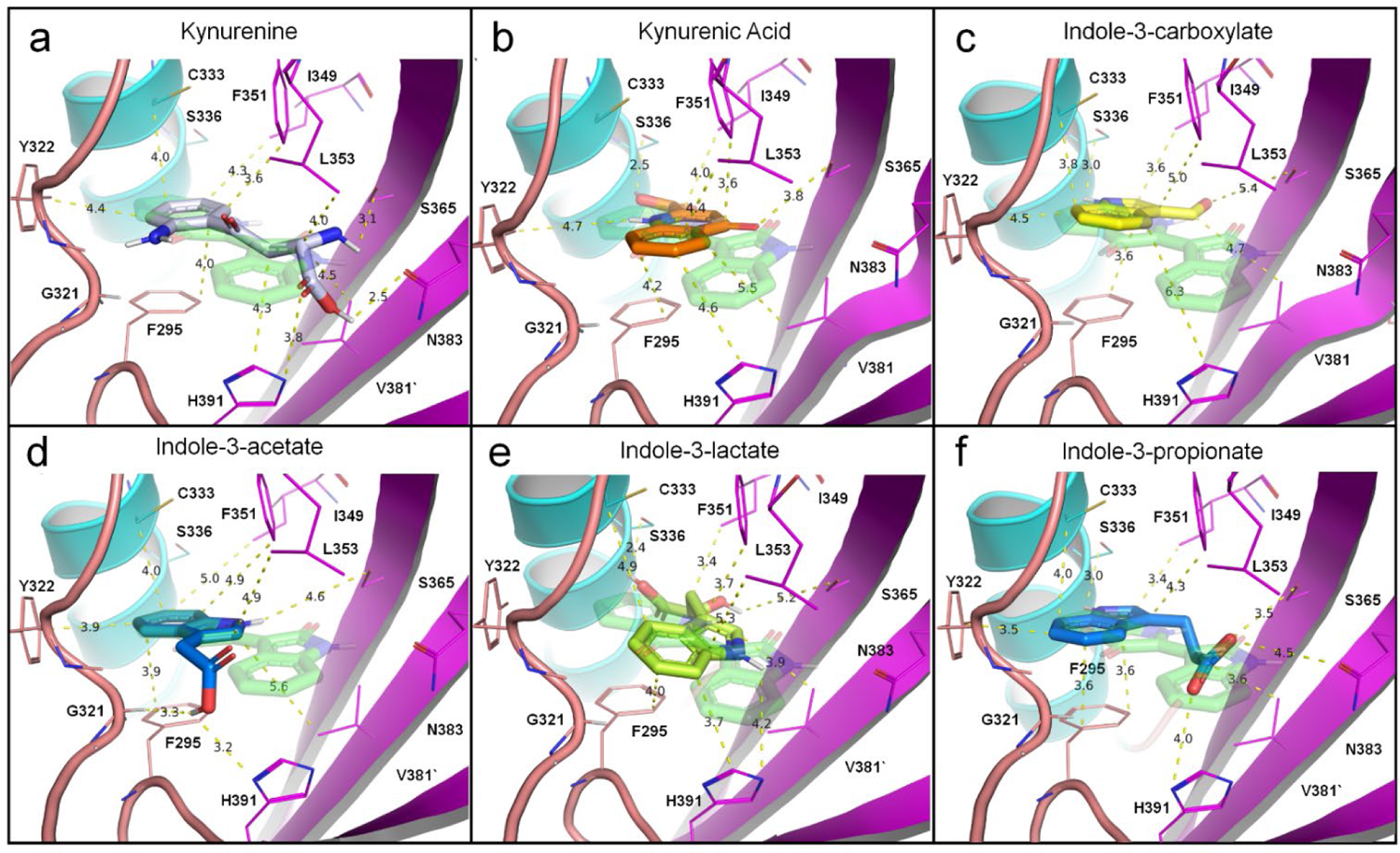
Trp metabolite docking in the Indirubin-bound model of the human AHR. As shown in Figure 1, the substrate binding properties of (a) Kynurenine; (b) Kynurenic acid, and four tryptophan-related metabolites, (c) Indole-3-carboxylate; (d) Indole-3-acetate; (e) Indole-3-lactate, and (f) Indole-3-propionate were explored using the cryo-EM structure of the AHR and Autodock Vina (see Table 1; see Methods). The results highlighted in Figure 1, are shown here with respect to the crystal structure coordinates of indirubin (shown in transparent green stick) in each panel, with respect to key active site residues: F295, G321, Y322, S336, S365, V381, N383 and H391. Key electrostatic and hydrophobic bond distances are shown (yellow dashes). High nanomolar affinity binding observed for indole-3-lactate, indole-3-propionate, and kynurenic acid suggest that a combination of discrete hydrophobic contacts (with F295, Y322, L349, F351, L353 and V381) and electrostatic contacts (with S336, S365, N383 and H391) are needed to fully engage the primary ligand binding domain of the human AHR.

**Table S1.**
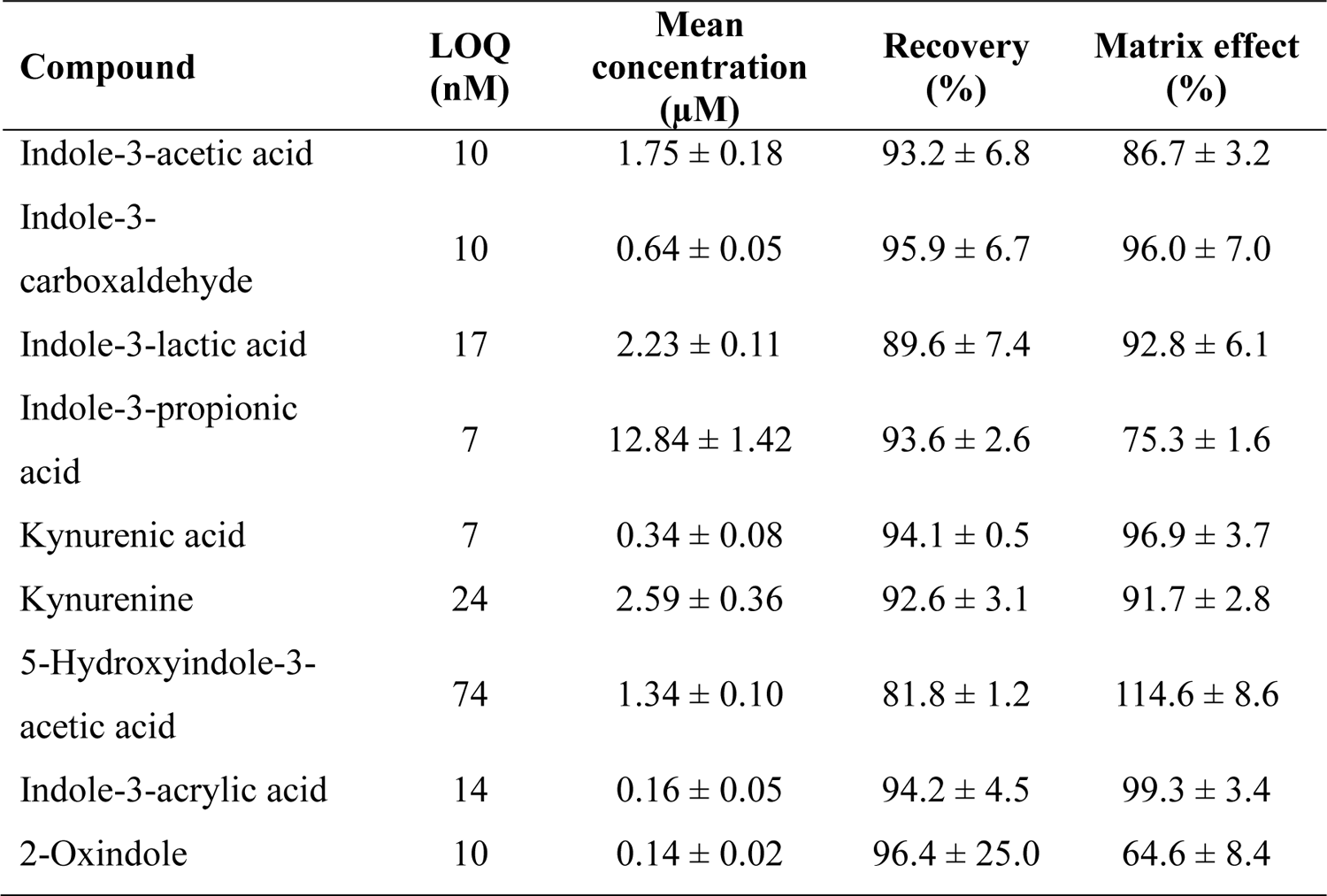
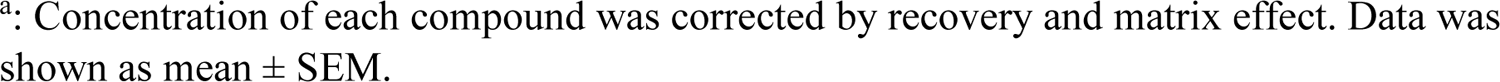
Parameters of AHR-related compounds quantified in serum of pathogen-free mice on a chow diet^a^..

| Compound | LOQ<br>(nM) | Mean<br>concentration<br>(μM) | Recovery<br>(%) | Matrix effect<br>(%) |
| --- | --- | --- | --- | --- |
| Indole-3-acetic acid | 10 | 1.75 ± 0.18 | 93.2 ± 6.8 | 86.7 ± 3.2 |
| Indole-3-carboxaldehyde | 10 | 0.64 ± 0.05 | 95.9 ± 6.7 | 96.0 ± 7.0 |
| Indole-3-lactic acid | 17 | 2.23 ± 0.11 | 89.6 ± 7.4 | 92.8 ± 6.1 |
| Indole-3-propionic acid | 7 | 12.84 ± 1.42 | 93.6 ± 2.6 | 75.3 ± 1.6 |
| Kynurenic acid | 7 | 0.34 ± 0.08 | 94.1 ± 0.5 | 96.9 ± 3.7 |
| Kynurenine | 24 | 2.59 ± 0.36 | 92.6 ± 3.1 | 91.7 ± 2.8 |
| 5-Hydroxyindole-3-acetic acid | 74 | 1.34 ± 0.10 | 81.8 ± 1.2 | 114.6 ± 8.6 |
| Indole-3-acrylic acid | 14 | 0.16 ± 0.05 | 94.2 ± 4.5 | 99.3 ± 3.4 |
| 2-Oxindole | 10 | 0.14 ± 0.02 | 96.4 ± 25.0 | 64.6 ± 8.4 |

<sup>a</sup>: Concentration of each compound was corrected by recovery and matrix effect. Data was shown as mean ± SEM.

**Table S2.** Parameters of AHR-related compounds quantified in human sera^a^.

| Compound | LOQ (nM) | Mean concentration (μM) | Recovery (%) | Matrix effect (%) |
| --- | --- | --- | --- | --- |
| Indole-3-acetic acid | 10 | 2.22 ± 0.10 | 92.5 ± 4.6 | 92.6 ± 3.6 |
| Indole-3-carboxaldehyde | 10 | 0.11 ± 0.01 | 73.0 ± 3.6 | 146.1 ± 18.4 |
| Indole-3-lactic acid | 17 | 1.92 ± 0.13 | 86.5 ± 8.5 | 85.2 ± 9.3 |
| Indole-3-propionic acid | 7 | 2.95 ± 0.27 | 98.5 ± 7.3 | 77.5 ± 7.0 |
| Kynurenic acid | 7 | 0.08 ± 0.01 | 80.0 ± 7.5 | 114.9 ± 8.3 |
| Kynurenine | 24 | 1.98 ± 0.11 | 80.2 ± 5.0 | 100.1 ± 4.0 |

<sup>a</sup>: Concentration of each compound was corrected by recovery and matrix effect. Data was shown as mean ± SEM.

**Table S3.**
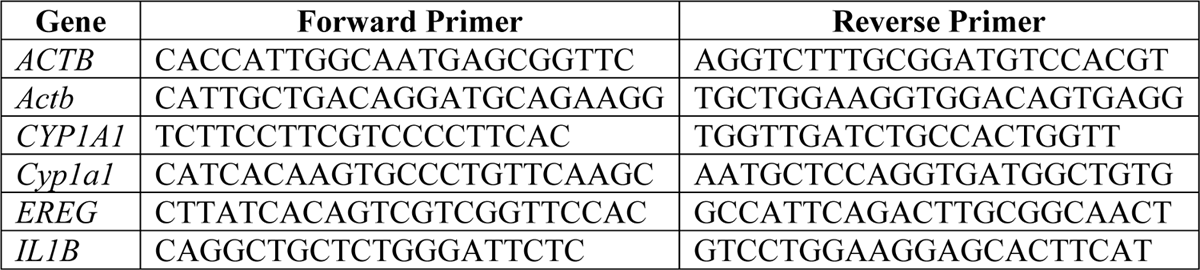
RT-qPCR primer sequences used in this study.

**Table S4.**
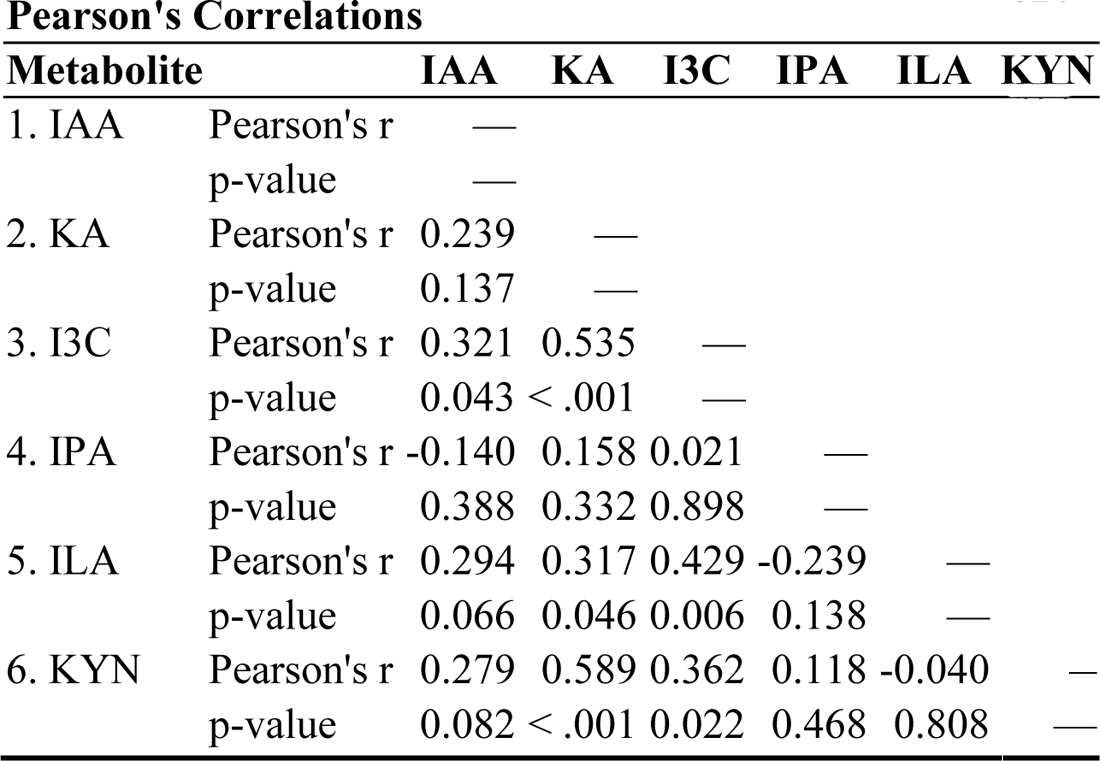
Pearson’s Correlations between tryptophan metabolites detected in human serum.

**Table S5.**
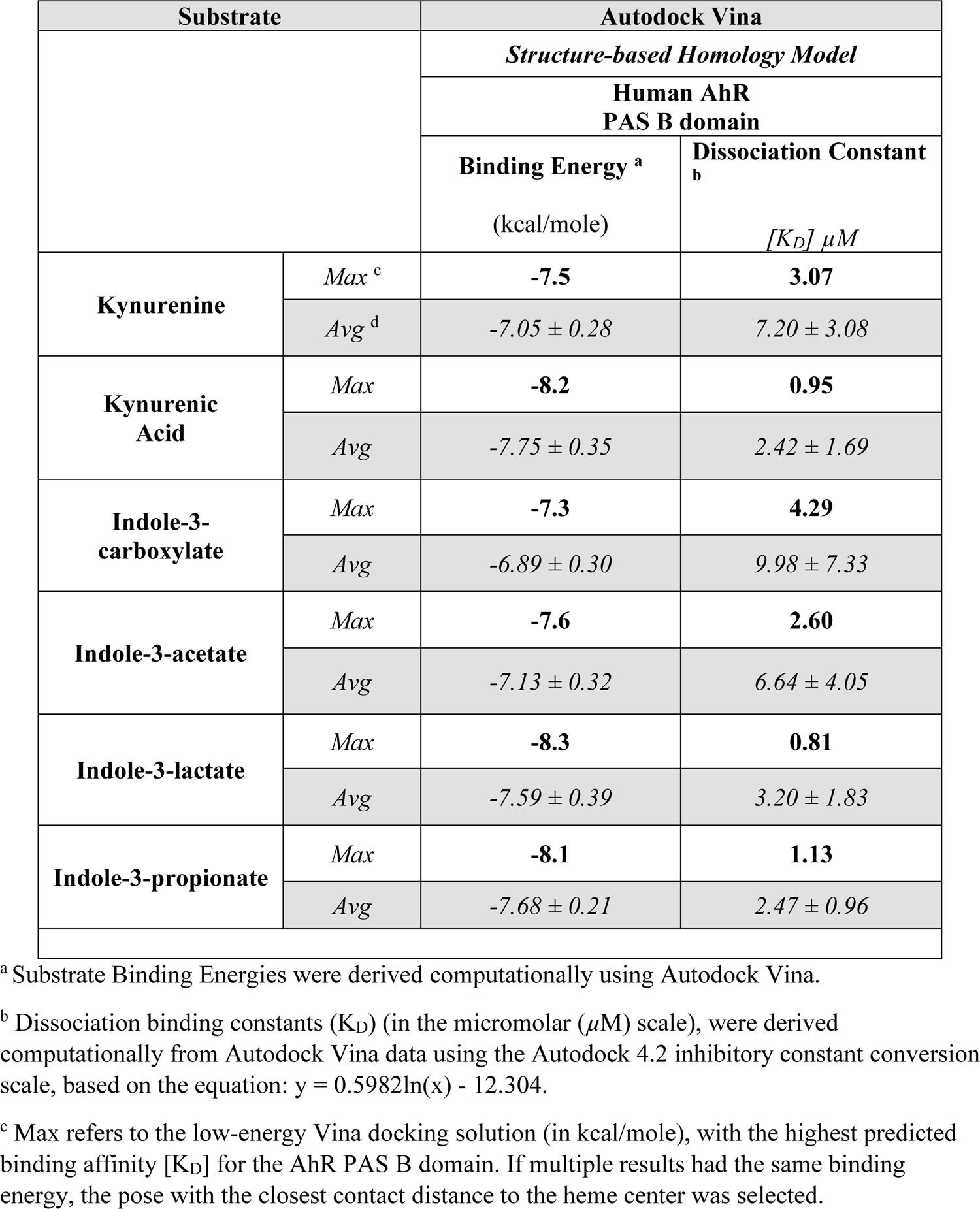
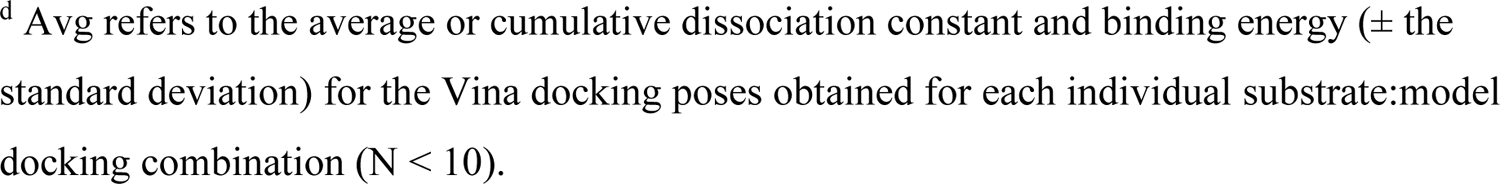
Computational Docking Analysis of Tryptophan Metabolites in the Human AHR PAS-B domain.

| Substrate |  | Autodock Vina |  |
| --- | --- | --- | --- |
|  |  | *Structure-based Homology Model* |  |
|  |  | Human AhR<br>PAS B domain |  |
|  |  | Binding Energy <sup>a</sup><br>(kcal/mole) | Dissociation Constant <sup>b</sup><br>[K<sub>D</sub>] μM |
| Kynurenine | Max <sup>c</sup> | -7.5 | 3.07 |
|  | Avg <sup>d</sup> | -7.05 ± 0.28 | 7.20 ± 3.08 |
| Kynurenic Acid | Max | -8.2 | 0.95 |
|  | Avg | -7.75 ± 0.35 | 2.42 ± 1.69 |
| Indole-3-carboxylate | Max | -7.3 | 4.29 |
|  | Avg | -6.89 ± 0.30 | 9.98 ± 7.33 |
| Indole-3-acetate | Max | -7.6 | 2.60 |
|  | Avg | -7.13 ± 0.32 | 6.64 ± 4.05 |
| Indole-3-lactate | Max | -8.3 | 0.81 |
|  | Avg | -7.59 ± 0.39 | 3.20 ± 1.83 |
| Indole-3-propionate | Max | -8.1 | 1.13 |
|  | Avg | -7.68 ± 0.21 | 2.47 ± 0.96 |

<sup>a</sup> Substrate Binding Energies were derived computationally using Autodock Vina.
<sup>b</sup> Dissociation binding constants (K<sub>D</sub>) (in the micromolar (μM) scale), were derived computationally from Autodock Vina data using the Autodock 4.2 inhibitory constant conversion scale, based on the equation: $y = 0.5982\ln(x) - 12.304$ .
<sup>c</sup> Max refers to the low-energy Vina docking solution (in kcal/mole), with the highest predicted binding affinity [K<sub>D</sub>] for the AhR PAS B domain. If multiple results had the same binding energy, the pose with the closest contact distance to the heme center was selected.
<sup>d</sup> Avg refers to the average or cumulative dissociation constant and binding energy ( ± the standard deviation) for the Vina docking poses obtained for each individual substrate:model docking combination ( $N < 10$ ).

